# Body temperature drives azole tolerance in *Candida albicans* by hindering the autophagic degradation of Erg11

**DOI:** 10.1101/2025.09.15.676225

**Authors:** Yanru Feng, Cheng Zhen, Wanqian Li, Malcolm Whiteway, Xinyu Fang, Xuqing Shen, Yuanying Jiang, Hui Lu

## Abstract

Human body temperature has been shown to limit the antifungal efficacy of azoles; however, the precise mechanisms underlying this limitation remain insufficiently understood. In this study, we observed that short-term exposure (24 h) to human body temperature (37□°C), in contrast to 30□°C, significantly enhances the tolerance of *Candida albicans* to azoles without inducing resistance. Our findings suggest that this increased tolerance is due to a reduction in the degradation of *C. albicans* Erg11 at 37□°C compared to 30□°C, thereby promoting azole tolerance. This phenomenon occurs because Erg11 degradation is mediated by autophagy, which is inhibited in *C. albicans* at 37□°C. The suppression of autophagy at this temperature may be associated with elevated mitochondrial production of reactive oxygen species, leading to mitochondrial dysfunction. In response to heat stress-induced mitochondrial dysfunction, autophagy-related proteins such as Atg8 accumulate around the mitochondria, while their levels in the endoplasmic reticulum decrease, consequently inhibiting autophagy in *C. albicans*. These findings suggest that promoting the degradation of Erg11 in *C. albicans* may serve as a viable strategy to augment the fungicidal efficacy of azoles in the human host.

## Introduction

In recent years, the morbidity and mortality rates associated with candidiasis have remained consistently high, presenting significant threats to public health and imposing a substantial medical burden(Lu et al., 2023a). Recent epidemiological data suggest that approximately 1.56 million individuals are affected by candidiasis annually, resulting in approximately 100,000 fatalities(Denning, 2024). Azoles have emerged as the most frequently utilized antifungal agents for the clinical prevention and treatment of fungal infections, owing to their broad antimicrobial spectrum, favorable safety profile, and diverse routes of administration(Shafiei et al., 2020). However, a primary challenge associated with azole therapy is their fungistatic nature, which allows *Candida albicans* to develop tolerance to azoles, ultimately leading to resistance(Berman and Krysan, 2020). Azole tolerance in *C. albicans* is demonstrated by its capacity to proliferate at azole concentrations exceeding the minimum inhibitory concentration (MIC) without an alteration in the azole MIC. In contrast, azole resistance is generally attributed to inherited mutations and is characterized by an increased azole MIC (Delarze and Sanglard, 2015; Feng et al., 2023; Xiong et al., 2025). Although the prevalence of fluconazole (FLC) resistance in *C. albicans* is generally low, reported at less than 1%(Pfaller et al., 2019), FLC often proves ineffective in treating candidiasis caused by susceptible *C. albicans* isolates. The observed discrepancy between overall treatment outcomes and minimal levels of clinical resistance is likely attributable to multiple factors(Lepak and Andes, 2014). Nevertheless, FLC tolerance in *C. albicans* is a critical consideration, as both in vitro and in vivo antifungal efficacy of FLC is substantially reduced against *C. albicans* strains exhibiting high azole tolerance(Li et al., 2024). Furthermore, elevated levels of FLC tolerance are associated with persistent candidemia(Rosenberg et al., 2018; Levinson et al., 2021), which contributes to approximately 50% of the mortality associated with neonatal candidiasis(Hammoud et al., 2013). However, the specific factors and mechanisms regulating the tolerance of *C. albicans* to azoles within the human host environment remain unclear.

It has been reported that the human body temperatures play a critical role in driving the development of drug resistance in pathogenic fungi. Studies have shown that, compared to a culture temperature of 30□°C, a temperature of 37□°C can decrease the susceptibility of *C. albicans* to azoles(Xu et al., 2021; Yang et al., 2023). Research on the fungal pathogen *Rhodosporidiobolus fluvialis* suggests that exposure to mammalian body temperature can induce mutagenesis, leading to the emergence of hypervirulent and pan-drug-resistant strains(Huang et al., 2024). Moreover, it has been demonstrated that anthropogenic global warming significantly affects azole resistance(Fisher et al., 2018; Fisher et al., 2022), with increased occurrences of azole-resistant *Aspergillus fumigatus* and *Candida parapsilosis* reported in warmer environments, such as greenhouses(Zhou et al., 2021) and tropical regions(Duong et al., 2021; Daneshnia et al., 2023). Additionally, the rapid global spread of multidrug-resistant *Candida auris* in humans has been linked to temperature adaptation driven by climate warming(Casadevall et al., 2019; Akinbobola et al., 2023). Body temperature may influence fungal drug susceptibility through non-genetic mechanisms within the human host(Bottery and Denning, 2024). However, the exact mechanisms by which short-term exposure (24 h) to body temperature reduces fungal susceptibility to azoles have yet to be elucidated. This gap in understanding poses a challenge for developing targeted antifungal strategies aimed at enhancing the efficacy of azoles in human hosts.

In this study, we observed that a short-term exposure to the human body temperature of 37□°C increases azole tolerance, but not azole resistance, in *C. albicans.* Our investigations further revealed that Erg11 is subject to degradation via the autophagy pathway. However, exposure to a body temperature of 37□°C, as opposed to 30□°C, inhibits autophagy, thereby decelerating the degradation rate of Erg11. This process ultimately enhances the tolerance of *C. albicans* to azoles. These findings expand the conceptual framework of azole tolerance mechanisms in *C. albicans*, demonstrating that the human host body temperature can modulate the degradation rate of the azole drug target Erg11 through the autophagy-dependent proteostasis regulation.

## Results

### Short-term exposure to body temperature enhances the tolerance of *C. albicans* to azoles

At a physiological body temperature of 37□°C, compared to 30□°C, we observed a decrease in the susceptibility of the *C. albicans* wild-type strain SN152 to FLC (Figure 1a). However, it remains uncertain whether this decreased drug susceptibility is due to an increase in the resistance level or tolerance level of *C. albicans* to FLC. In this study, we characterized the enhancement of FLC resistance in *C. albicans* by assessing the elevation of the minimum inhibitory concentration (MIC) value of FLC, while the increase in the super-MIC growth (SMG) value was used to represent the elevation of FLC tolerance in *C. albicans*(Rosenberg et al., 2018; Lu et al., 2023b) (Figure 1b). Our findings indicate that, after a short-term exposure, compared to 30□°C, the temperature of 37□°C significantly increased the SMG value of FLC against the strain SN152 from 0.070 to 0.440, whereas the MIC value of FLC against the strain SN152 remained constant at 0.5 μg/ml (Figure 1c). Cyclosporin A, a known calcineurin inhibitor, was observed to abolish the tolerance of *C. albicans* to FLC by reducing the SMG value of FLC, but not affect the resistance of *C. albicans* to FLC, as indicated by the unchanged MIC value(Rosenberg et al., 2018; Lu et al., 2023b). Our study demonstrated that the decreased susceptibility of *C. albicans* to FLC after incubation at 37□°C for 24 h was reversed by cyclosporin A (Extended Data Figure 1a). Furthermore, the deletion of the *CDR1* gene, which encodes the efflux pump Cdr1, resulted in a reduced MIC value of FLC against *C. albicans*, yet it did not eliminate FLC tolerance in the organism(Rosenberg et al., 2018; Wang et al., 2022). Our findings indicated that, in comparison to 30□°C, culturing at 37□°C further diminished the susceptibility of the *C. albicans cdr1*Δ/Δ null mutant to FLC (Extended Data Figure 1b). This observation is attributed to the physiological temperature, which increased the SMG value of FLC against the *cdr1*Δ/Δ null mutant, although it did not elevate the MIC value for this mutant (Figure 1c). These results suggest that the reduced susceptibility of *C. albicans* to FLC following short-term exposure to physiological temperature is primarily due to an enhanced tolerance, rather than the development of resistance of *C. albicans* to azoles.

**Figure 1.**
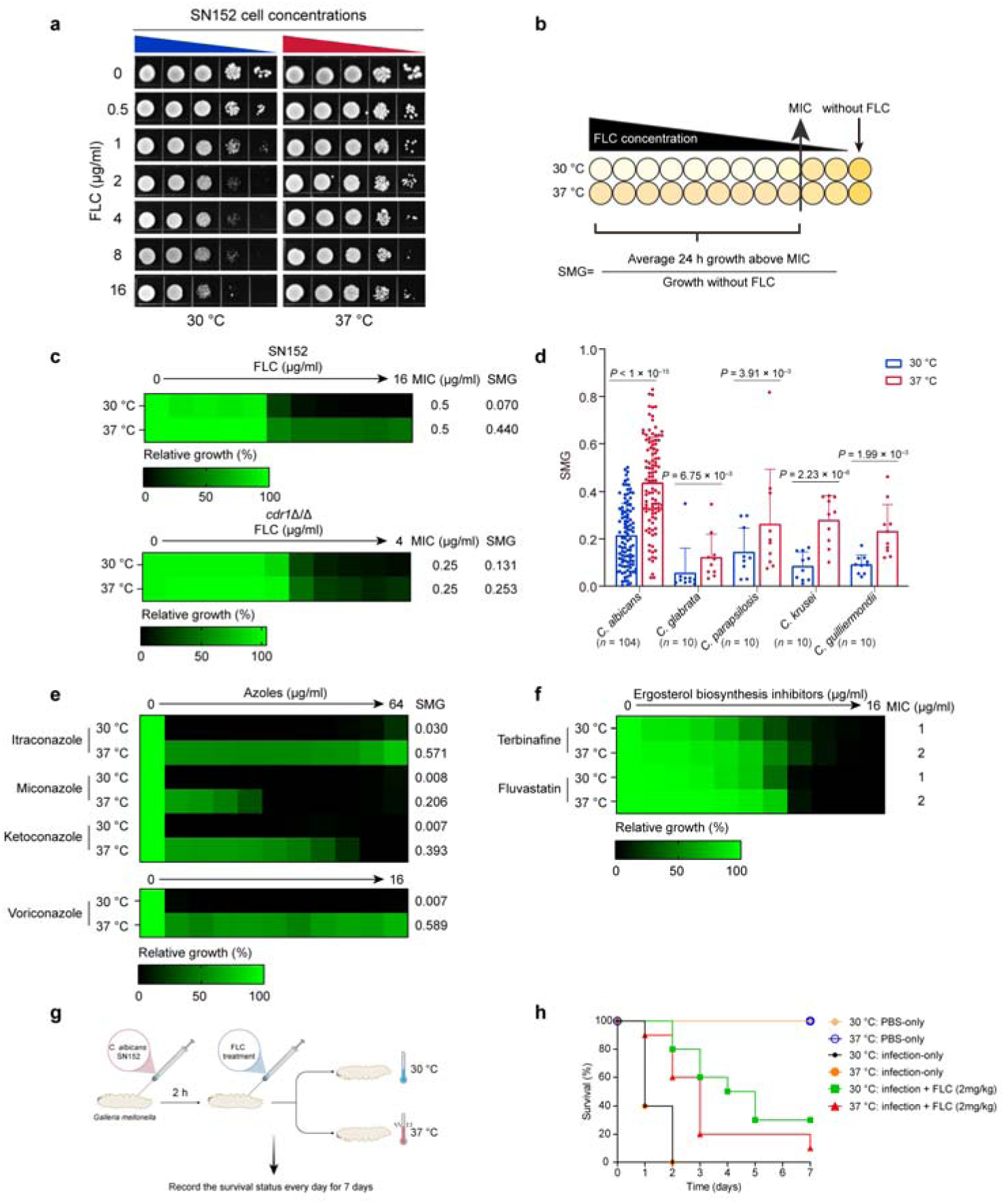
Short-term exposure to body temperature enhances the tolerance of *C. albicans* to azoles. **(a)** Spotting assays of *C. albicans* SN152. Cells were spotted onto YPD agar containing different concentrations (0 μg/ml, 0.5 μg/ml, 1 μg/ml, 2 μg/ml, 4 μg/ml, 8 μg/ml and 16 μg/ml) of fluconazole (FLC) and cultured at 30□°C or 37□°C for 2□days. The different colors of right triangle indicate cells were 1:10 serially diluted and cultured at 30□°C (blue one) or 37□°C (red one). (**b**) Illustration of minimum inhibitory concentration (MIC) and supra-MIC growth (SMG) calculations. MIC was calculated at 24 h as the FLC concentration at which 50% of the growth was inhibited at 30□°C, relative to growth without FLC. SMG was calculated as the average growth per well above the MIC divided by the level of growth without FLC. The darker the yellow color, the more vigorous the cell growth. **c**, Broth microdilution assays showing MIC and SMG levels at 24 h for *C. albicans* wild-type SN152 and *cdr1*Δ/Δ null mutant cultured at 30□°C or 37□°C in the presence of FLC. **d**, The values of SMG of FLC against clinical *Candida* isolates which were cultured at 30□°C or 37□°C. Data represent mean□±□standard deviation (s.d). The *P* values were determined by the Wilcoxon matched-pairs signed rank test for *C. albicans* (*n* = 104) and *C. parapsilosis* (*n* = 10) and the two-tailed paired *t* test for *C. glabrata* (*n* = 10), *C. krusei* (*n* = 10) and *C. guilliermondii* (*n* = 10). **e**, Broth microdilution assays showing SMG levels at 24 h for *C. albicans* SN152 cultured at 30□°C or 37□°C in the presence of azoles. Technical duplicates were averaged. **f**, Broth microdilution assays showing MIC levels at 24 h for *C. albicans* SN152 cultured at 30□°C or 37□°C in the presence of terbinafine and fluvastatin. Technical duplicates were averaged. **g**, Schematic of *C. albicans* SN152 infection and FLC treament models in *Galleria mellonella* larvae. *G. mellonella* larvae were infected with *C. albicans* SN152, and the FLC treatment was administered after 2 hour-infection with a single dose. All groups were incubated at 30 °C or 37 °C, whose survival status was recorded daily during the 7-day experimental period. **h**, The survival curves of *G. mellonella* larvae (*n* = 10 per group) over 7 days. Larvae were subjected to three treatment conditions: PBS□only controls (uninfected, no FLC treatment), infection with *C. albicans* SN152 alone, or infection with *C. albicans* SN152 plus 2□mg/kg FLC. Each condition was incubated at either 30□°C or 37□°C. Survival was assessed daily.

Our investigation revealed that elevating the culture temperature from 30□°C to 37□°C significantly increased the SMG values of FLC against clinical isolates of *C. albicans* (*n* = 104), *Candida glabrata* (*n* = 10), *Candida parapsilosis* (*n* = 10), *Candida krusei* (*n* = 10), and *Candida guilliermondii* (*n* = 10) (Figure 1d). This observation suggests that physiological body temperature generally enhances the tolerance of *Candida* species isolates to FLC. Additionally, the SMG values for itraconazole, miconazole, ketoconazole and voriconazole against *C. albicans* were significantly elevated when cells were cultured at 37□°C (Figure 1e). Moreover, the antifungal efficacy of other ergosterol biosynthesis inhibitors, such as terbinafine (targeting Erg1) and fluvastatin (targeting Hmg1), were reduced at 37□°C (Figure 1f). However, the culture temperature did not affect the antifungal activity of amphotericin B (binding and destroying plasma ergosterol), caspofungin (targeting Fks1 to inhibit the synthesis of 1,3-β-D-glucan), tunicamycin (inducing ER stress), myriocin (blocking serine-palmitoyltransferase in sphingolipid biosynthesis) and staurosporine (inhibiting protein kinase C) against *C. albican*s (Extended Data Figure 1c). Overall, these findings indicate that brief exposure to physiological body temperature substantially increases the tolerance of *C. albicans* to azoles. Additionally, to further verify that body temperature diminishes the therapeutic effect of FLC, we chose the poikilotherm, *Galleria mellonella (G. mellonella)*, to construct an experimental model, which has been reported can grow normally at both 30 °C(Brennan et al., 2002) and 37 °C(Fuchs et al., 2010; Marena et al., 2025). The wild-type *C. albicans* strain SN152 was used to infect *G.mellonella* larvae and FLC treatment was administered after 2 h-infection with a single dose, then larvae were incubated at 30 °C or 37 °C for the duration of the experiment (Figure 1g). Over a 7-day experimental period, the survival rates of *G. mellonella* in the 2 mg/kg FLC treatment groups at 30□°C (median survival = 4.5 days) were higher than that in the corresponding groups at 37□°C (median survival = 3 days) (Figure 1h). These in vivo data demonstrate that the elevated temperature compromised the antifungal effect of FLC.

### The augmented tolerance of *C. albicans* to azoles at body temperature is dependent on Erg11

Upon short-term exposure to body temperature, *C. albicans* exhibited reduced susceptibility to inhibitors targeting ergosterol biosynthesis, including azoles, terbinafine and fluvastatin (Figures 1e and 1f). Conversely, this exposure did not significantly impact the efficacy of inhibitors targeting non-ergosterol biosynthesis pathways, such as caspofungin, tunicamycin, myriocin, and staurosporine (Extended Data Figure 1c). Furthermore, our study demonstrated that both fluvastatin (1 μg/ml) and terbinafine (1 μg/ml) effectively decreased the SMG values of FLC against *C. albicans* at 37□°C (Figure 2a). These findings suggest that exposure to body temperature may influence the ergosterol biosynthesis pathway in *C. albicans*, thereby enhancing its tolerance to azoles.

**Figure 2.**
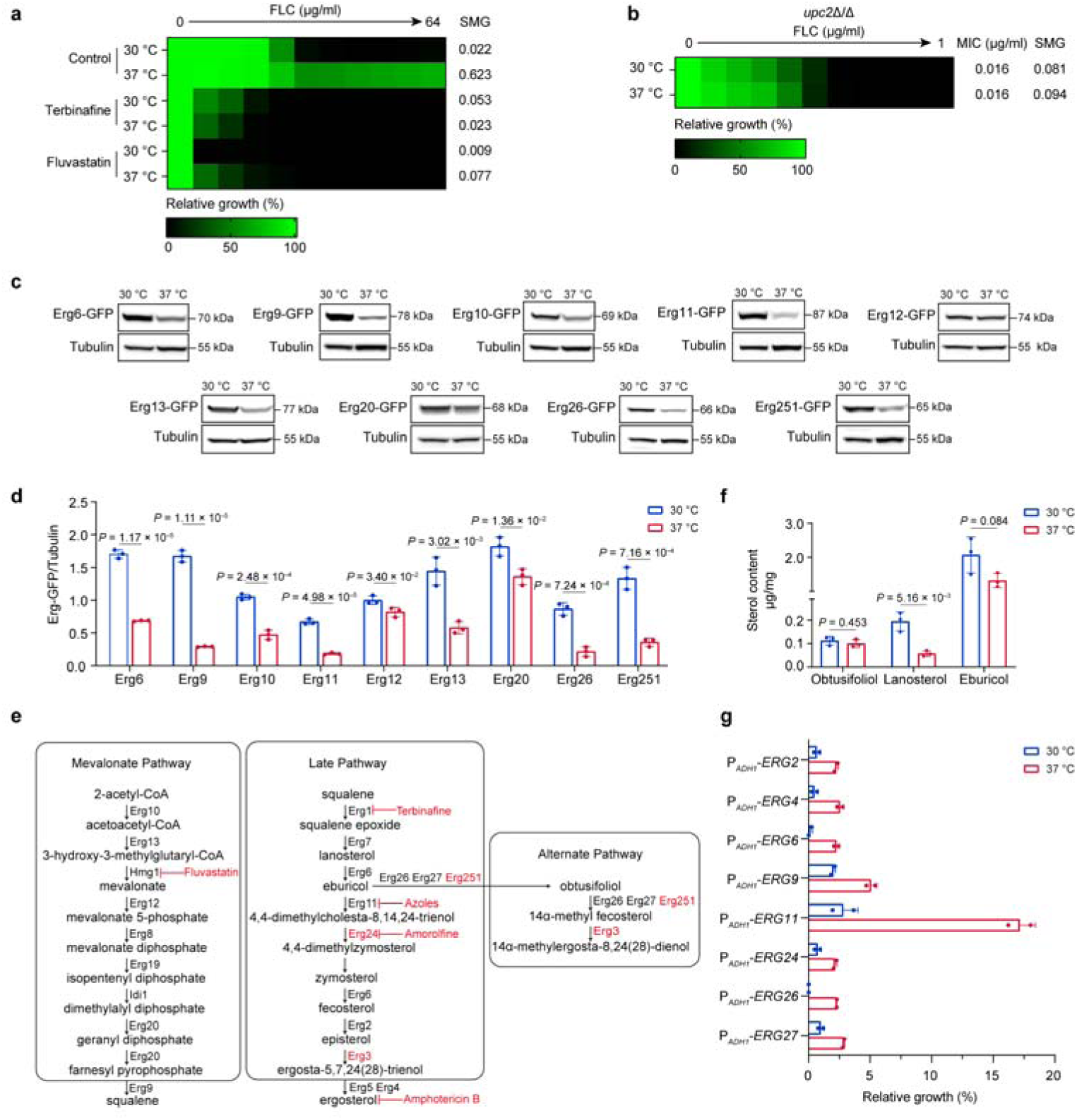
Body temperature enhances *C. albicans*’ tolerance to azoles through Erg11. **a**, Broth microdilution assays showing SMG levels at 24 h for *C. albicans* SN152 cultured at 30□°C or 37□°C in the presence of FLC combined with fluvastatin (1 μg/ml) or terbinafine (1 μg/ml). The control group represents cells were in the presence of FLC without combined compounds. **b**, Broth microdilution assays showing MIC and SMG levels at 24 h for *upc2*Δ/Δ null mutant cultured at 30□°C or 37□°C in the presence of FLC. Technical duplicates were averaged. **c**, The expression of proteins involved in ergosterol synthesis (Erg6, Erg9, Erg10, Erg11, Erg12, Erg 13, Erg20, Erg26 and Erg251) examined by western blotting. Cells were precultured at 30□°C or 37□°C for 16 h. The representative images shown are from three replicates. **d**, Quantification of the signal intensity ratio of Erg-GFP to Tubulin by Image J. Data represent means□±□s.d. of three replicates, two-tailed, unpaired t test. **e**, Illustration of ergosterol synthesis pathway and ergosterol synthesis inhibitors. **f**, The determination of sterol content in the presence of 16 μg/ml FLC based on GC-MS at the 16-hour time point. Data represent means□±□s.d. of three replicates, two-tailed, unpaired t test. **g**, The relative growth at optical density at 600 nm (OD_600_) of *upc2*Δ/Δ null mutants in which the expression of ergosterol biosynthesis genes (*ERG2*, *ERG4*, *ERG6, ERG9*, *ERG11*, *ERG24, ERG26*, and *ERG27*) were increased by ADH1 promoter in the presence of 0.25 μg/ml FLC at 30□°C or 37□°C for 24 h. Technical duplicates were averaged.

We proceeded to delete the *EFG1*, *NDT80*, and *UPC2* genes in *C. albicans*, which encode transcriptional regulators involved in ergosterol biosynthesis(Lo et al., 2005; Sellam et al., 2009; Vasicek et al., 2014; Yang et al., 2015). Our findings demonstrated that at 37□°C, there was an increase in the SMG value of FLC against the *efg1*Δ/Δ and *ndt80*Δ/Δ null mutants (Extended Data Figure 2a). In contrast, the *upc2*Δ/Δ null mutant consistently exhibited low MIC and SMG values for FLC at both 30□°C and 37□°C (Figure 2b). This observation was further corroborated by spot assays, which confirmed that 37□°C did not reduce the susceptibility of the *upc2*Δ/Δ null mutant to FLC (Extended Data Figure 2b). These results suggest that Upc2 may contribute to the tolerance of *C. albicans* to azoles at physiological body temperature.

Body temperature may activate Upc2 transcription, thereby enhancing azole tolerance in *C. albicans*. We tagged the C-terminal of Upc2 with GFP, and subsequent laser confocal microscopy demonstrated that brief exposure to body temperature did not influence the nuclear import of Upc2 in *C. albicans* in the presence of FLC (16 μg/ml) (Extended Data Figures 2c and 2d). Furthermore, RNA-seq analysis indicated that 4 h of incubation at 37□□ did not significantly alter the transcription levels of most ergosterol synthesis genes (Extended Data Figure 2e). Additionally, we tagged the C-terminal regions of key enzymes involved in ergosterol biosynthesis with GFP, including Erg6, Erg9, Erg10, Erg11, Erg12, Erg13, Erg20, Erg26, and Erg251. Consistent with the RNA-seq findings, no upregulation of these enzymes was observed at 37□°C compared to 30□°C. In fact, western blot analysis suggested a slight decrease in expression levels at 37□°C (Figures 2c and 2d). Furthermore, the analysis conducted using gas chromatography-mass spectrometry (GC-MS) indicated that there were no notable differences in intracellular ergosterol concentrations between the two temperature conditions at the 4-hour, 8-hour, and 16-hour incubation intervals (Extended Data Figure 2f). These results collectively imply that the enhanced tolerance of *C. albicans* to azoles at 37□°C is not mediated by the activation of Upc2 transcriptional activity.

In *C. albicans*, Erg7 (lanosterol synthase) catalyzes the conversion of squalene epoxide to lanosterol. This lanosterol is subsequently transformed into eburicol through the activity of Erg6 (Δ-(24)-sterol C-methyltransferase). Following this conversion, Erg11 mediates the oxidative demethylation of the 14-α-methyl group from eburicol, producing 4,4-dimethylcholesta-8,14,24-trienol(Martel et al., 2010). Inhibition of Erg11 by azoles induces an alternative biosynthetic pathway involving Erg251 (C-4 sterol methyl oxidase), Erg26 (C-3 sterol dehydrogenase), Erg27 (3-keto sterol reductase), and Erg3 (C-5 sterol desaturase), which results in the synthesis of compensatory sterols that retain 14α-methyl groups(Lu et al., 2023b) (Figure 2e). GC-MS analysis demonstrated a significant reduction in lanosterol levels, with no statistically significant changes in obtusifoliol concentrations, when cells were exposed to 16 μg/ml FLC at 37□°C compared to 30□°C (Figure 2f). Furthermore, utilizing the *ADH1* promoter to enhance the expression of ergosterol biosynthesis genes in the *upc2*Δ/Δ null mutant, specifically *ERG2*, *ERG4*, *ERG6, ERG9*, *ERG11*, *ERG24, ERG26*, and *ERG27*, showed that only the upregulation of *ERG11* significantly improved the relative growth of *C. albicans* under FLC exposure (0.25 μg/ml) at 37□°C (Figure 2g). These results are corroborated by chemical genomics experiments, which demonstrated that fluvastatin, targeting Hmg1, and terbinafine, targeting Erg1 — both acting upstream of Erg11 — can inhibit azole tolerance induced by heat stress (Figure 2a). Conversely, amorolfine, which targets Erg24 and is positioned downstream of Erg11(Polak-Wyss et al., 1985), did not show a similar inhibitory effect (Extended Data Figure 2g). Additionally, a notable increase in FLC tolerance was observed in the *erg24*Δ/Δ null mutant at 37□°C (Extended Data Figure 2h). The deletion of either the *ERG251* or *ERG3* gene(Lu et al., 2023b), which are components of an alternative pathway, did not abolish the heat stress-induced tolerance to azoles (Extended Data Figure 2i). These findings suggest that body temperature augments the tolerance of *C. albicans* to azoles via a mechanism reliant on Erg11.

### Body temperature inhibits the degradation of Erg11

We further modulated the expression of the *ERG11* and *ENO1* genes in the *ERG11*/*erg11*Δ and *ENO1*/*eno1*Δ heterozygous deletion mutants using the doxycycline (DOX)-inducible Tet-off promoter system (Extended Data Figure 3a). This system facilitates the regulation of the transcription levels of the *ERG11* and *ENO1* genes in response to varying DOX concentrations (Extended Data Figure 3b). Notably, the MIC values of DOX against the P_tet-off_-*ENO1*/*eno1*Δ mutant remained consistent at both 30□°C and 37□°C. Conversely, the MIC value of DOX against the P_tet-off_-*ERG11*/*erg11*Δ mutant at 37□°C was observed to be twice as high as at 30□°C (Figure 3a). These findings indicate that the increased tolerance of *C. albicans* to FLC at 37□°C cannot be attributed to the expression level of Erg11 at this temperature.

**Figure 3.**
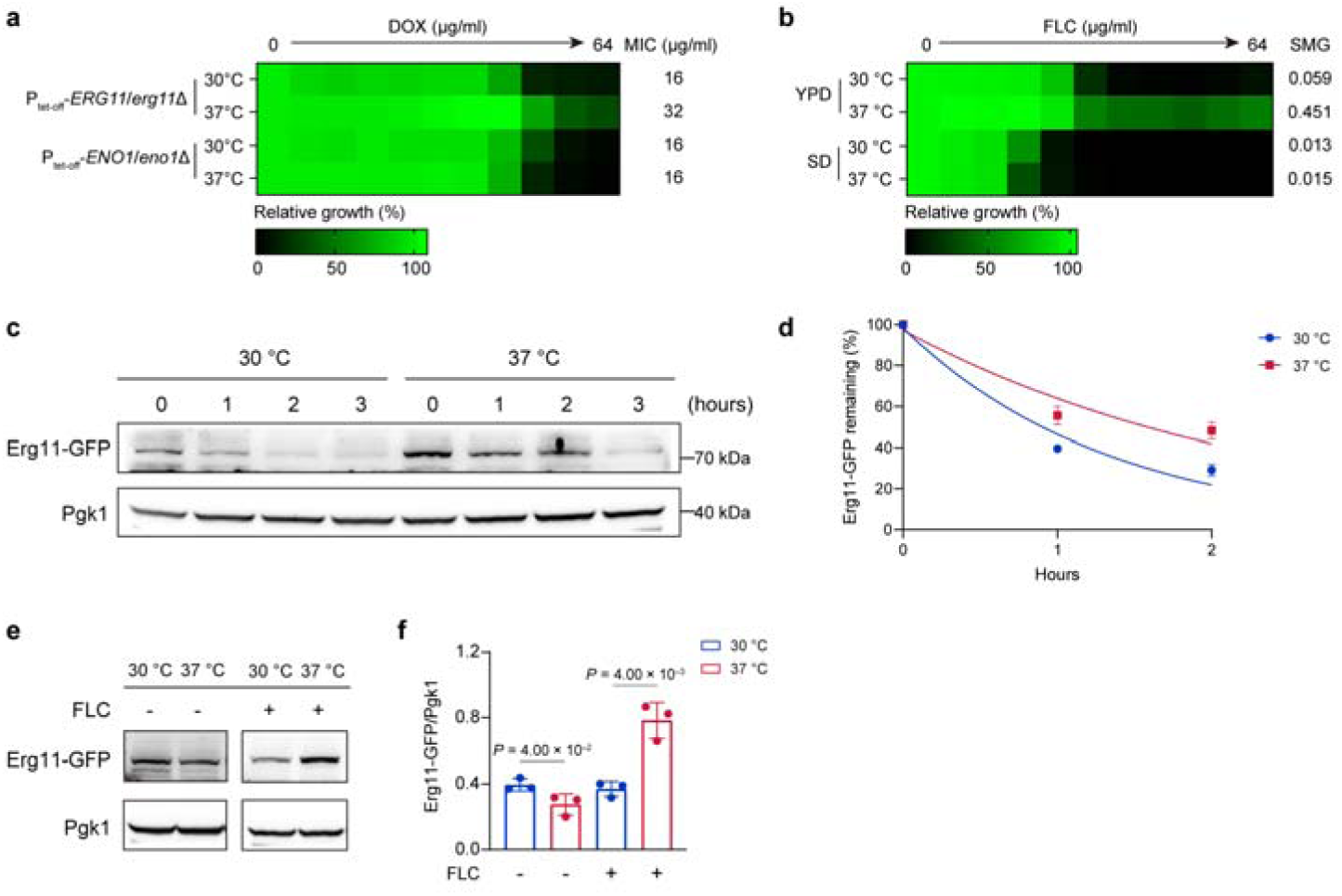
Body temperature inhibits the degradation of Erg11. **a**, Broth microdilution assays showing MIC levels at 24 h for P_tet-off_-*ERG11*/*erg11*Δ mutant and P_tet-off_-*ENO1*/*eno1*Δ mutant cultured at 30□°C or 37□°C in the presence of DOX. Technical duplicates were averaged. **b**, Broth microdilution assays showing SMG levels at 24 h for *C. albicans* SN152 cultured in different culture medium (YPD or SD) at 30□°C or 37□°C in the presence of FLC. Technical duplicates were averaged. **c**, Degradation of Erg11-GFP in *C. albicans* SN152 after inhibition of protein synthesis by cycloheximide upon a 3-hour shift to 30 °C or 37 °C. The representative images shown are from three replicates. **d**, Quantification of the degradation of Erg11-GFP calculated as the ratio of Erg11-GFP to Pgk1 by Image J. Set the protein level at the initial time point to 100 %. Data represent means□±□s.d. of three replicates, two-tailed, unpaired t test. The decay curves were fitted using a one-phase exponential decay model (*Y* = *Y*_0_ • *e*^−*Kt*^) based on all replicate data points. **e**, The expression of Erg11 protein examined by western blotting. Cells were precultured at 30□°C and 37□°C for 4 h with or without 4 μg/ml FLC. The representative images shown are from three replicates. **f**, Quantification of the signal intensity ratio of Erg11-GFP to Pgk1 by Image J. Data represent means□±□s.d. of three replicates, two-tailed, unpaired t test.

Moreover, the induction of azole tolerance by body temperature was determined to be contingent upon the culture medium. Specifically, an increased azole tolerance phenotype at 37□°C was observed in the nutrient-rich YPD medium, whereas this was not the case in the nutrient-poor SD medium (Figure 3b). These results imply that the heightened tolerance of *C. albicans* to FLC at 37□°C cannot be ascribed to an augmented catalytic activity of Erg11 at this temperature.

Cycloheximide (CHX) chase analysis was employed to assess the degradation rates of Erg11 at 30□°C and 37□°C. The analysis revealed that, unlike at 30□°C, the degradation of Erg11 was markedly inhibited at 37□°C (Figure 3c). The decay curves were fitted to a one-phase exponential decay model. At 30□°C, Erg11 exhibited a relatively short half-life of approximately 0.93 h (95% CI: 0.74 to 1.17 h, *R*^2^=0.96). In contrast, at 37□°C, the degradation rate was markedly slower, with an estimated half-life of 1.65 h (95% CI: 1.25 to 2.33 h, *R*^2^=0.91) (Figure 3d). At 37□°C, the inhibition of Erg11 degradation leads to the sustained preservation of its catalytic activity. As a result, *C. albicans* requires a reduced amount of Erg11 to produce an equivalent amount of ergosterol at 37□°C compared to 30□°C. This induces a sterol-mediated negative feedback mechanism that decreases Erg11 levels in *C. albicans* at 37□°C in the absence of FLC (Figures 3e and 3f). However, when FLC inhibits Erg11 activity, the resulting reduction in ergosterol content triggers an upregulation of Erg11 expression. Given that 37□°C suppresses Erg11 degradation, the exposure of azoles leads to elevated levels of Erg11 protein in *C. albicans* at 37□°C relative to 30□°C (Figures 3e and 3f), thereby enhancing the azole tolerance of *C. albicans*.

### Erg11 undergoes degradation through autophagy

In *S. cerevisiae,* it has been documented that Erg11 is subject to degradation via the Asi-mediated Endoplasmic Reticulum-Associated Degradation (ERAD) pathway following ubiquitination (Foresti et al., 2014; Khmelinskii et al., 2014; Natarajan et al., 2020). However, the precise mechanism by which ubiquitinated Erg11 is degraded in *C. albicans*—whether through the 26S proteasome or autophagy in *C. albicans—*remains unclear. To investigate the degradation pathway of Erg11 in *C. albicans*, we employed wortmannin, an autophagy inhibitor(Delorme-Axford et al., 2015), and MG132, a proteasome inhibitor(Marshall and Vierstra, 2022), to ascertain their effects on Erg11 degradation. Our results demonstrated that wortmannin significantly enhanced the FLC tolerance of *C. albicans* at 30□°C and further augmented FLC tolerance at 37□°C. Conversely, MG132 diminished FLC resistance at both 30□°C and 37□°C, and reduced FLC tolerance at 37□°C (Figure 4a). Moreover, CHX chase analysis revealed that wortmannin, but not MG132, inhibited the degradation of Erg11 at 30□°C (Figures 4b and 4c). To further explore the key *ATG* genes associated with FLC tolerance in *C. albicans*, we deleted the core ATG genes (*ATG1*, *ATG3*, *ATG7*, *ATG8*, *ATG9*, *ATG13* and *ATG18*). Spot assay results showed that only the *atg8*Δ/Δ null mutants exhibited increased FLC tolerance at 30 °C (Figure 4d). Additionally, CHX chase analysis revealed that, in contrast to the wild-type strain SN152, deletion of the *ATG8* gene inhibited Erg11 degradation (Figure 4e and 4f). These results confirm that Erg11 degradation is associated with autophagy pathway, particularly dependent on the *ATG8* gene. We employed the CellTracker Blue CMAC dye to stain vacuoles and tagged Erg11 with GFP. Laser confocal microscopy experiments revealed that Erg11 was localized within vacuoles, as indicated by the co-localization of green and red fluorescence, resulting in a yellow fluorescence signal (Figures 4g and 4h). Furthermore, our findings revealed that incubation at 37□°C, as opposed to 30□°C, significantly hindered the entry of Erg11 into vacuoles (Figures 4g and 4h). These results collectively suggest that Erg11 undergoes degradation via autophagy, and that this process may be inhibited at 37□°C, thereby suppressing the degradation of Erg11.

**Figure 4.**
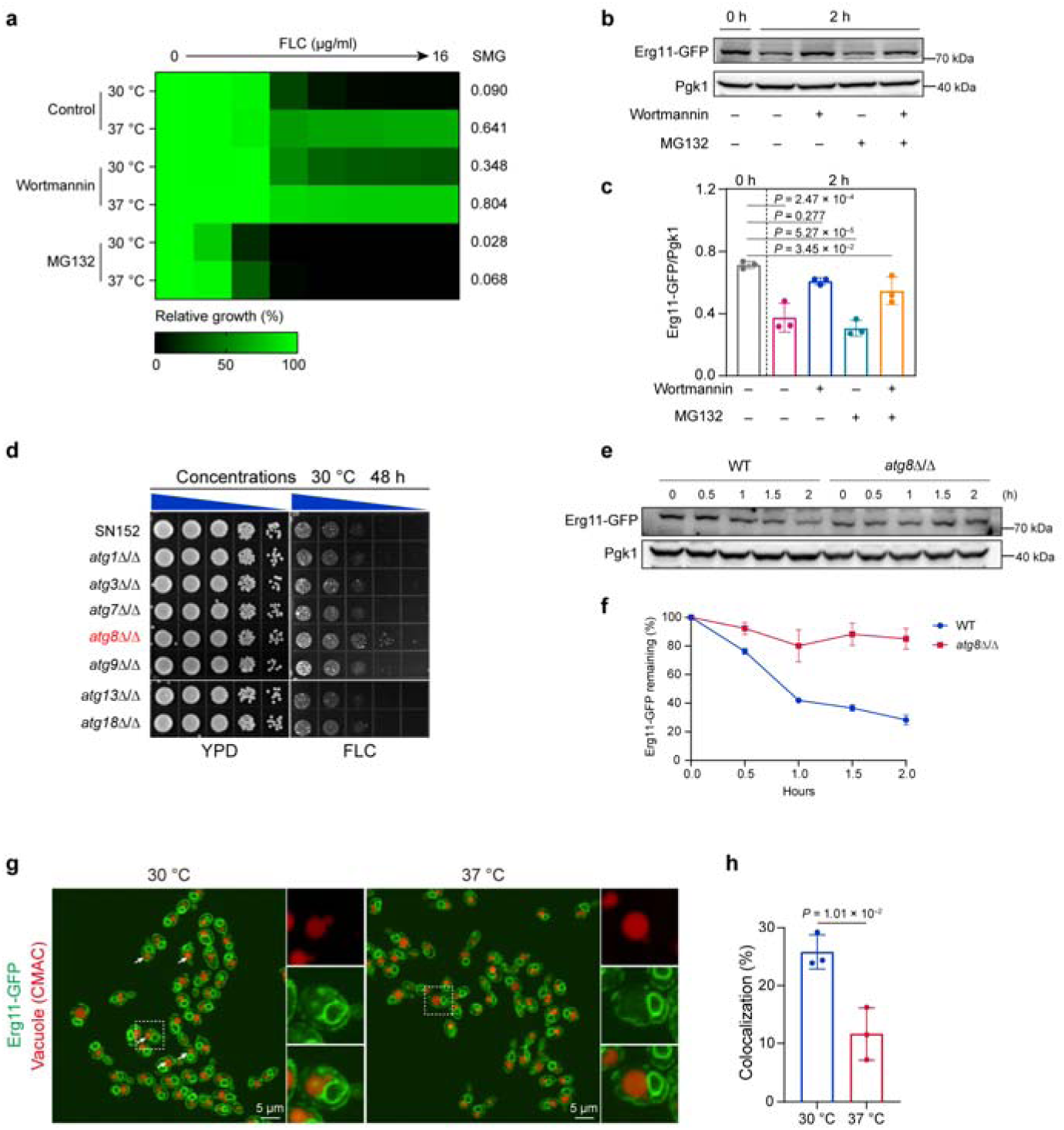
Erg11 undergoes degradation through autophagy. **a**, Broth microdilution assays showing SMG levels at 24 h for *C. albicans* SN152 cultured at 30□°C or 37□°C in the presence of FLC combined with wortmannin (1 μM) or MG132 (6.25 μM). Technical duplicates were averaged. **b**, Degradation of Erg11-GFP in *C. albicans* SN152 in the presence of Wortmannin (1 μM) and MG132 (100 μM) after inhibition of protein synthesis by cycloheximide upon a 2-hour shift to 30□°C. **c**, Quantification of the degradation of Erg11-GFP calculated as the ratio of Erg11-GFP to Pgk1 by Image J. Data represent means□±□s.d. of three replicates, ordinary one-way ANOVA, Bonferroni’s multiple comparisons test. **d**, Spotting assays of *C. albicans* SN152, *atg1*Δ/Δ, *atg3*Δ/Δ, *atg7*Δ/Δ, *atg8*Δ/Δ, *atg9*Δ/Δ, *atg13*Δ/Δ, *atg18*Δ/Δ null mutants. Cells were spotted onto YPD agar with or without 4 μg/ml of fluconazole (FLC) and cultured at 30□°C for 2□days. The blue right triangle indicate cells were 1:10 serially diluted and cultured at 30□°C. **e**, Degradation of Erg11-GFP in *C. albicans* SN152 and *atg8*Δ/Δ null mutant after inhibition of protein synthesis by cycloheximide upon a 2-hour shift to 30 °C. The representative images shown are from three replicates. **f**, Quantification of the degradation of Erg11-GFP calculated as the ratio of Erg11-GFP to Pgk1 by Image J. Set the protein level at the initial time point to 100 %. Data represent means□±□s.d. of three replicates, two-tailed, unpaired t test. **g**, Colocalization between Erg11-GFP and vacuole in cells precultured at 30□°C or 37□°C for 1 h. Pseudo-color was used to display the blue fluorescence of the vacuole dye (CellTracker Blue CMAC) as red. The white arrow indicated vacuoles with Erg11-GFP. The representative images shown are from three replicates. Scale bar, 5□μm. **h**, Quantification of the colocalization ratio of the number of cells colocalized with Erg11 and vacuoles to the total number of cells. The data represent means□±□s.d of three replicates, two-tailed, unpaired t test.

### Body temperature inhibits autophagy in *C. albicans*

Considering that body temperature inhibits the degradation of Erg11, through the autophagy pathway, it can be posited that body temperature may suppress the autophagic process in *C. albicans*. To investigate this hypothesis, we employed a *C. albicans* GFP-Atg8 mutant, in which GFP was fused to the N-terminus of Atg8(Zhen et al., 2024b). During the activation of autophagy, GFP-Atg8 localizes to autophagosomes and is subsequently released into vacuoles. Notably, GFP exhibits greater resistance to vacuolar hydrolases, leading to the degradation of Atg8 while GFP remains intact. Thus, the conversion of GFP-Atg8 to GFP serves as an indicator of autophagic activity. Our findings indicated that at 37□°C, the ratio of GFP to GFP-Atg8 was significantly lower compared to that observed at 30□°C (Figures 5a and 5b).

**Figure 5.**
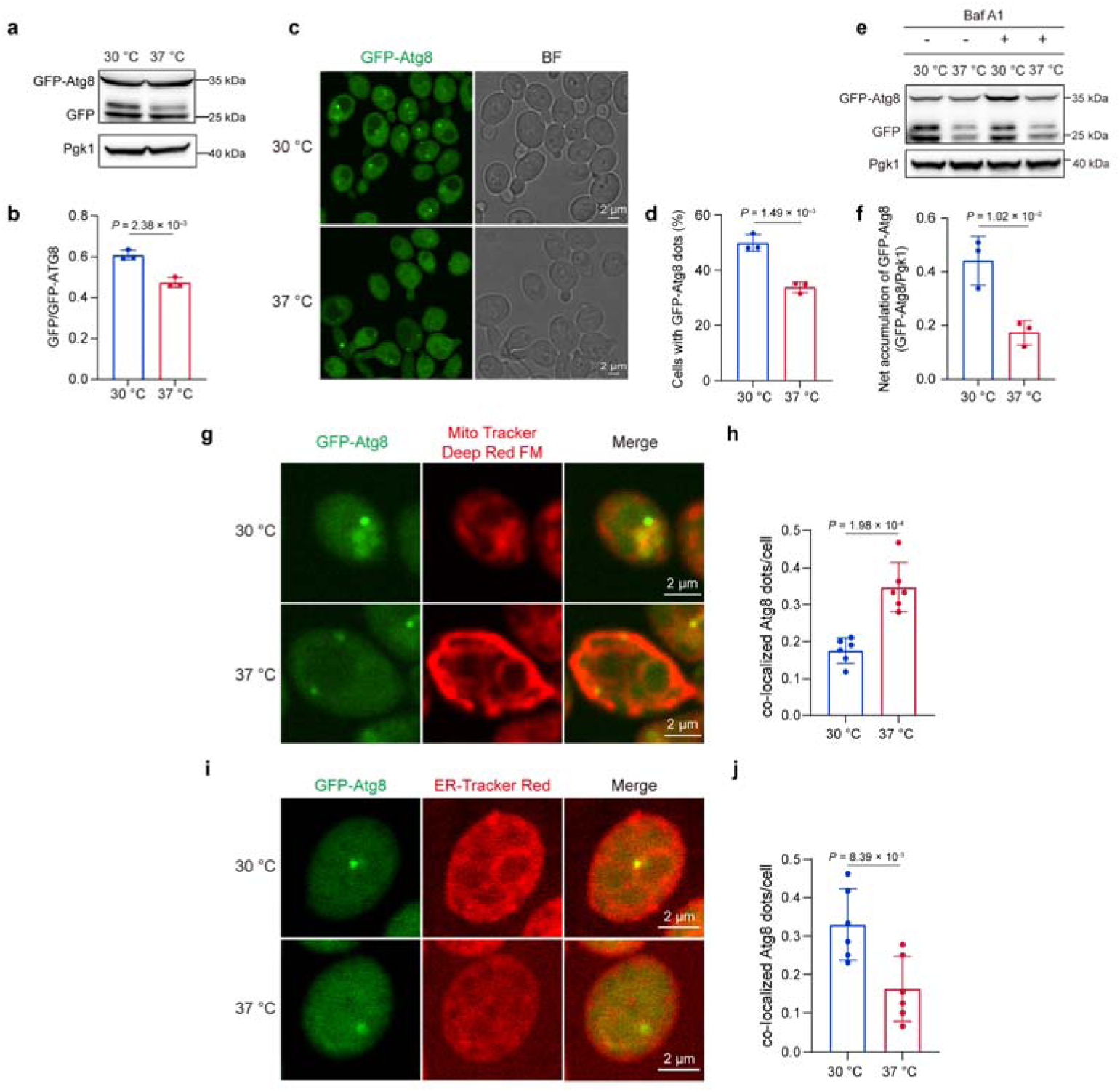
Body temperature inhibits autophagy in *C. albicans*. **a**, GFP-ATG8 cleavage assay showing the autophagic activity after pre-cultivation at 30 °C or 37 °C for 4 h. **b**, Quantification of the autophagic activity displaying as the ratio of free GFP to GFP-ATG8 by Image J. Data represent means□±□s.d. of three replicates, two-tailed, unpaired t test. **c**, Confocal images of GFP-ATG8 dots in cells precultured at 30 °C or 37 °C for 1 h. BF represents cells in bright field. Scale bar, 2□μm. **d**, The ratio of the number of cells with GFP-ATG8 dots to the total number of cells. Data represent means□±□s.d. form three replicates, two-tailed, unpaired t test. **e**, Bafilomycin A1 inhibition assays showing the autophagic flux in *C. albicans* expressing GFP-Atg8 after pre-cultivation at 30 °C or 37 °C for 4 h. Baf A1 (2 μM) or vehicle (DMSO) was added at 2 h post□incubation. Pgk1 served as loading control. **f**, Quantification of the net accumulation of full-length GFP-Atg8 calculating as the normalized band intensity of Baf A1□treated samples minus that of vehicle□treated samples (Δ = (GFP-Atg8/Pgk1)_+_Baf A1 − (GFP-Atg8/Pgk1)□Baf A1). Data represent means□±□s.d. of three replicates, two-tailed, unpaired t test. **g**, Colocalization between GFP-Atg8 dots and mitochondria in cells precultured at 30□°C or 37□°C for 30 min. Scale bar, 2□μm. **h**, Quantification of the colocalization ratio of the number of GFP-Atg8 dots colocalized with mitochondria to the number of cells containing GFP-Atg8 dots. The data represent means□±□s.d. n□=□6 confocal microscopic images form three replicates, two-tailed, unpaired t test. **i**, Colocalization between GFP-Atg8 dots and ER in cells precultured at 30□°C or 37□°C for 30 min. Scale bar, 2□μm. **j**, Quantification of the colocalization ratio of the number of GFP-Atg8 dots colocalized with ER to the number of cells containing GFP-Atg8 dots. The data represent means□±□s.d. n□=□6 confocal microscopic images form three replicates, two-tailed, unpaired t test.

Additionally, laser confocal microscopy demonstrated that at 30□°C, *C. albicans* cells exhibited a higher number of cells with green fluorescent dots, which are indicative of autophagosomes containing GFP-Atg8; conversely, at 37□°C, the number of such cells was markedly diminished (Figures 5c and 5d). In order to quantitatively measure the autophagic flux, Bafilomycin A1 (Baf A1) inhibition assays combined with Western blot analysis of GFP-Atg8 were conducted. Baf A1 is a specific inhibitor of vacuolar□type ATPase (V□ATPase), which blocks autophagosome-lysosome fusion, while leaving autophagosome formation unaffected(Chang et al., 2020). The autophagic flux was calculated as the difference in full-length GFP-Atg8 levels (normalized to loading control, Pgk1) between BafA1□treated and vehicle□treated (without BafA1) samples (Δ = Baf A1-vehicle), which reflects the net accumulation of autophagosomes during the treatment period(Metur and Klionsky, 2024; Paumier et al., 2025). The results showed that in both temperature conditions, Baf A1 blocked autophagosome-lysosome fusion, resulting in further accumulation of GFP-Atg8 compared to the group without Baf A1 (Figure 5e). The net accumulation (Δ) of full-length GFP-Atg8 was significantly higher in 30 °C than in 37 °C (Δ_30 °C = 0.443 ± 0.075, Δ_37 °C = 0.174 ± 0.036, *P* = 1.02 × 10^−2^ ), indicating a higher autophagosome formation rate at 30 °C (Figure 5f). These results confirm that 37 °C decreased autophagosome formation to reduce complete autophagic flux. In addition, we utilized MitoTracker Deep Red FM dye for mitochondrial labeling, ER-Tracker Red dye for ER labeling, and GFP for Atg8 labeling, employing confocal laser scanning microscopy to examine the effects of elevated environmental temperature on the intracellular localization of Atg8. Our findings indicate that, relative to the condition at 30□°C, the proximity of Atg8 to mitochondria increased at 37□°C (Figures 5g and 5h), whereas its colocalization with the ER diminished (Figures 5i and 5j), consequently resulting in the inhibition of autophagy.

### Body temperature-induced mitochondrial oxidative damage that inhibits autophagy

Research has demonstrated that heat stress induces mitochondrial dysfunction via oxidative stress mechanisms(Chang-Chien et al., 2024). In this study, we cultured *C. albicans* at 30□°C and 37□°C for 4 h, followed by RNA sequencing analysis. Principal component analysis (PCA) revealed clear separation between the two groups (Extended Data Figure 4a). Using thresholds of |log2FC| ≥ 1 and Q value < 0.05, we identified a total of 1,733 differentially expressed genes (DEGs), including 932 significantly up-regulated and 801 down-regulated genes (Figure 6a). A heatmap of the top 20 DEGs ranked by Q-value further confirmed distinct expression profiles between cells grown at 37 °C versus 30 °C (Extended Data Figure 4b).

**Figure 6.**
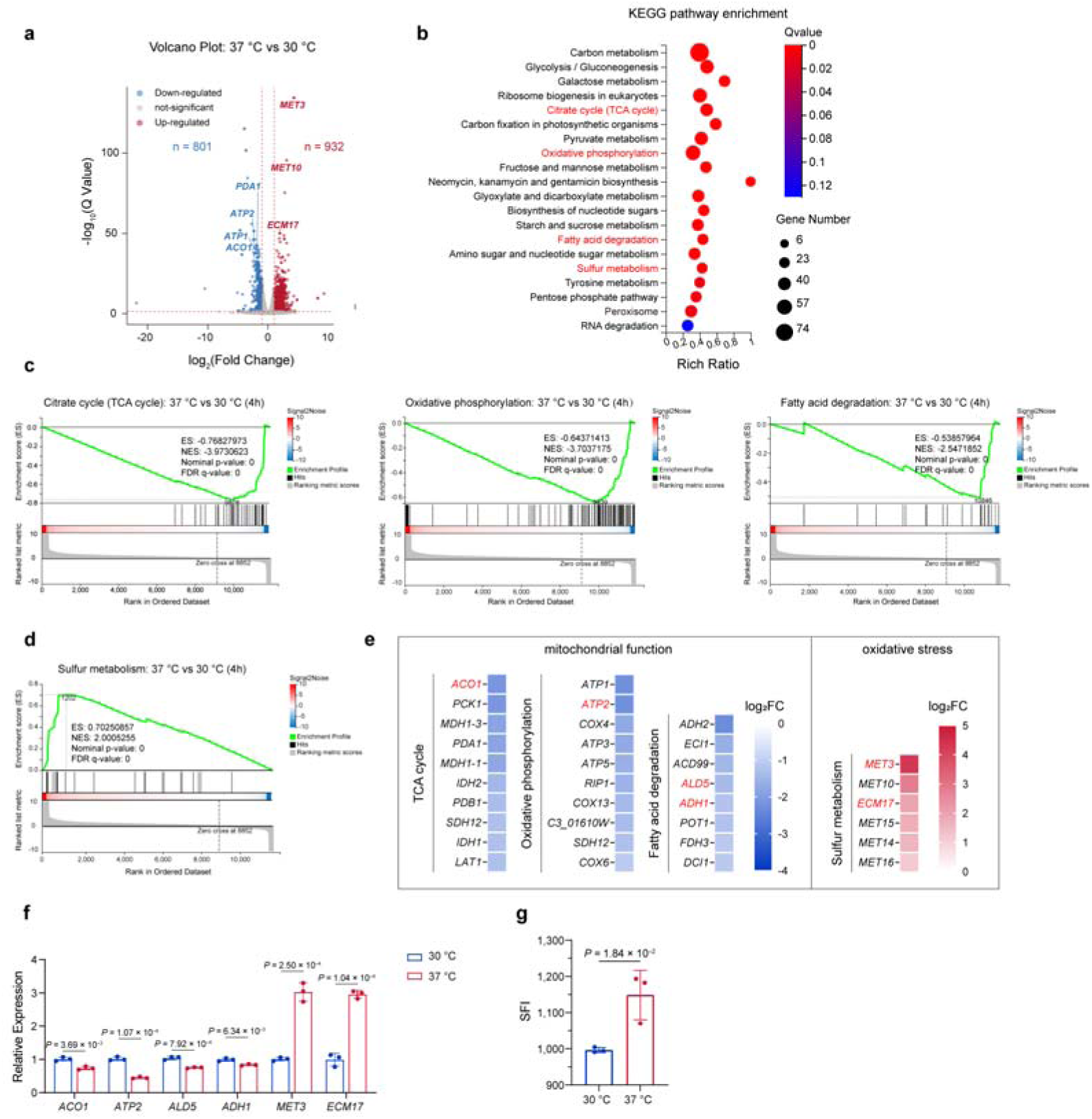
Body temperature induced the mitochondrial oxidative damage. **a**, Volcano plot of transcriptomics data shows differentially expressed genes (DEGs) (cut-off: |log2FC| ≥ 1 and Q-value < 0.05 indicated as red dashed line) between 30 °C and 37 °C cultured strains after 4-h incubation. **b**, Kyoto Encyclopedia of Genes and Genomes (KEGG) enrichment analysis for DEGs between 30 °C and 37 °C cultured strains after 4-h incubation. The Rich Ratio is calculated as the ratio of differentially expressed genes with in a pathway term to all annotated genes within that pathway term. **c**, Gene set enrichment analysis (GSEA) shows downregulation of Citrate cycle (TCA cycle), Oxidative phosphorylation and Fatty acid degradation for strains cultured at 37 °C. Enrichment Scores (ES) indicate downregulated genes in strains cultured at 37 °C. **d**, GSEA shows upregulation of Sulfur metabolism for strains cultured at 37 °C. Enrichment Scores (ES) indicate upregulated genes in strains cultured at 37 °C. **e**, Heatmap of the top DEGs in TCA cycle, Oxidative phosphorylation, Fatty acid degradation and Sulfur metabolism pathways based on GSEA, ordered by |log2FC|. **f**, mRNA levels of genes (*ACO1*, *ATP2*, *ALD5*, *ADH1*, *MET3* and *ECM17*) related to mitochondrial function or oxidative stress measured with qRT–PCR in *C. albicans* SN152 precultured at 30 °C or 37 °C for 4 h. The signal was normalized to the strains cultured at 30 °C. Data represent means□±□s.d. of three replicates, two-tailed, unpaired t test. **g**, The intracellular ROS levels measured by specific fluorescence intensity (SFI) of the oxidant-sensitive probe 2′,7′-dichlorofluorescin diacetate (DCFH-DA) in cells precultured at 30□°C or 37□°C for 30 min. SFI was calculated as fluorescence [(Em = 535 nm/Ex = 485 nm)/mg protein]. Data represent means□±□s.d. of three replicates, two-tailed, unpaired t test.

To characterize the biological processes affected by elevated temperature, we performed Kyoto Encyclopedia of Genes and Genomes (KEGG) enrichment and Gene Ontology (GO) analyses on the identified DEGs. KEGG enrichment showed significant enrichment of the citrate cycle (TCA cycle), oxidative phosphorylation and fatty acid degradation pathways (Figure 6b). These pathways are central to mitochondrial energy metabolism: the TCA cycle generates reducing equivalents (Yang et al., 2024), fatty acid degradation supplies acetyl-CoA (Enkler et al., 2023), and oxidative phosphorylation drives ATP production (Ni and Gao, 2025; Yang et al., 2025). Their enrichment among the DEGs indicates that elevated temperature broadly impacts mitochondrial function. Additionally, the sulfur metabolism pathway was significantly enriched, which is linked to glutathione biosynthesis and cellular antioxidant defense (Park et al., 2024; Shangguan et al., 2024).

Consistent with these findings, GO enrichment analysis, including cellular component (CC), biological process (BP), and molecular function (MF), revealed that the most significantly enriched terms were associated with mitochondrial energy metabolism and stress responses (Extended Data Figure 4c-4e). Together, these transcriptomic profiles suggest that exposure to 37 °C not only impairs mitochondrial energy metabolism but also triggers an oxidative stress response in *C. albicans*.

To further assess pathway-wide coordinate regulation, we performed Gene Set Enrichment Analysis (GSEA) using the full ranked gene list. GSEA demonstrated significant and coordinated downregulation of the TCA cycle (NES = -3.97, FDR q-value < 0.05), oxidative phosphorylation (NES = -3.70, FDR q-value < 0.05), and fatty acid degradation (NES = -2.55, FDR q-value < 0.05) pathways at 37 °C (Figure 6c). This global transcriptional suppression strongly indicates widespread impairment of mitochondrial function. Conversely, the sulfur metabolism pathway was significantly upregulated (NES = +2.00, FDR q-value < 0.05) (Figure 6d). In *C. albicans*, sulfur metabolism pathway supports the biosynthesis of sulfur-containing amino acids, which serve as precursors for glutathione—a major cellular antioxidant. Activation of this pathway thus represents a classic adaptive response to oxidative stress.

We next highlighted the top DEGs within these pathways based on Q-value with |log2FC| ≥ 1 (Figure 6e) and selected representative core genes with known roles in mitochondrial function or oxidative stress for qRT-PCR validation. Expression levels of the *ACO1* ( encoding aconitase, a key enzyme in the TCA cycle) (Farooq et al., 2013), *ATP2* ( encoding the F1 beta subunit of F1F0 ATPase complex)(Li et al., 2018), *ALD5* ( encoding a mitochondrial aldehyde dehydrogenase (NAD□) ) and *ADH1* genes (encoding the fermentative alcohol dehydrogenase)(Francis et al., 2007) were significantly decreased at 37 °C compared to 30 °C (Figure 6f). In contrast, the *MET3* ( encoding ATP sulfurylase)(Park et al., 2024) and *ECM17* genes (encoding the beta subunit of sulfite reductase) were significantly up-regulated (Figure 6f).

To further investigate, we employed 2′,7′-dichlorofluorescin diacetate (DCFH-DA) to quantify reactive oxygen species (ROS) levels and observed that the 37□°C condition, compared to 30□°C, resulted in elevated ROS levels in *C. albicans* (Figure 6g). These results demonstrate that the elevated temperature induced the mitochondrial oxidative damage.

An antioxidant Diphenyleneiodonium chloride (DPI), which selectively inhibits intracellular ROS, was shown to effectively synergize with FLC to mitigate the tolerance of *C. albicans* at 37□°C (Figure 7a), implying that mitochondrial oxidative damage in *C. albicans* induced by body temperature may contribute to the increased tolerance of *C. albicans* to azoles. Moreover, the inhibition intracellular ROS by DPI increased the autophagic activity at 37 °C, exhibiting that the higher ratio of GFP to GFP-Atg8 in DPI-treated group (Figure 7b and 7c). Additionally, the degradation rate of Erg11 was accelerated by the treatment of the antioxidant DPI at 37 °C (Figure 7d and 7e). These findings indicated that the clearance of ROS can increased the autophagic activity and shorten the half-life of Erg11, exhibiting the synergistic effect with FLC. Under conditions of heat stress, Atg8 in plants has been documented to engage biological processes beyond autophagy, including its relocation to the Golgi apparatus to facilitate Golgi reorganization(Zhou et al., 2023). Additionally, based on the results that the deletion of *ATG8* increased the FLC tolerance of *C. albicans* at 30□°C and prolonged the half-life of Erg11 (Figure 4e and 4f) and the proximity of Atg8 to mitochondria increased at 37□°C (Figures 5g and 5h), whereas its colocalization with the ER diminished (Figures 5i and 5j), we hypothesized that the redistribution of Atg8 may provide protection to mitochondria against oxidative damage induced by body temperatures. To test this hypothesis, we used the *atg8*Δ/Δ null mutant in *C. albicans* and assessed the growth rates of both the *atg8*Δ/Δ null mutant and its parental strain SN152 at 30□°C and 37□°C, respectively. Our findings indicated that the *atg8*Δ/Δ null mutant exhibited growth rates comparable to those of the SN152 strain at 30□°C. However, at 37□°C, the *atg8*Δ/Δ null mutant demonstrated reduced growth relative to the SN152 strain (Figure 7f), suggesting that Atg8 plays a critical role in the response to oxidative stress at 37□°C. Additionally, we employed JC-1 to evaluate the impact of the *ATG8* gene on mitochondrial membrane potential (MMP). The lipophilic JC-1 dye, with a delocalized positive charge, accumulates in functional mitochondria, emitting red fluorescence. In dysfunctional mitochondria, it remains dispersed in the cytosol, emitting green fluorescence. A decrease in red and increase in green fluorescence signals mitochondrial dysfunction. Mitochondria with a red-to-green ratio over 1 are considered normal (P1 in Figure 7g), while those below 1 are dysfunctional (P2 in Figure 7g). Our findings indicated that at 30□°C, the SN152 strain (0.290% ± 0.096%) and the *atg8*Δ/Δ null mutant (0.730% ± 0.131%) exhibit comparable proportions of cells in the P2 region (*P* > 0.999). In contrast, at 37□°C, the *atg8*Δ/Δ null mutant (8.42% ± 0.800%) shows a significantly higher proportion of cells in the P2 region compared to the SN152 strain (5.33% ± 1.19%; *P* = 1.07 × 10^−2^) (Figures 7g and 7h). Additionally, the proportion of cells in the P2 region at 37 °C was significantly higher for both the SN152 strain (*P* = 4.52 × 10^−4^ ) and the *atg8*Δ/Δ null mutant (*P* = 2.08 × 10^−5^) than that at 30 °C, showing that the elevated temperatures resulted in mitochondrial dysfunction. These results suggest that mitochondrial dysfunction in *C. albicans*, triggered by elevated temperatures, may lead to a reduced presence of autophagy-related proteins, such as Atg8, within the ER, thereby inhibiting autophagy.

**Figure 7.**
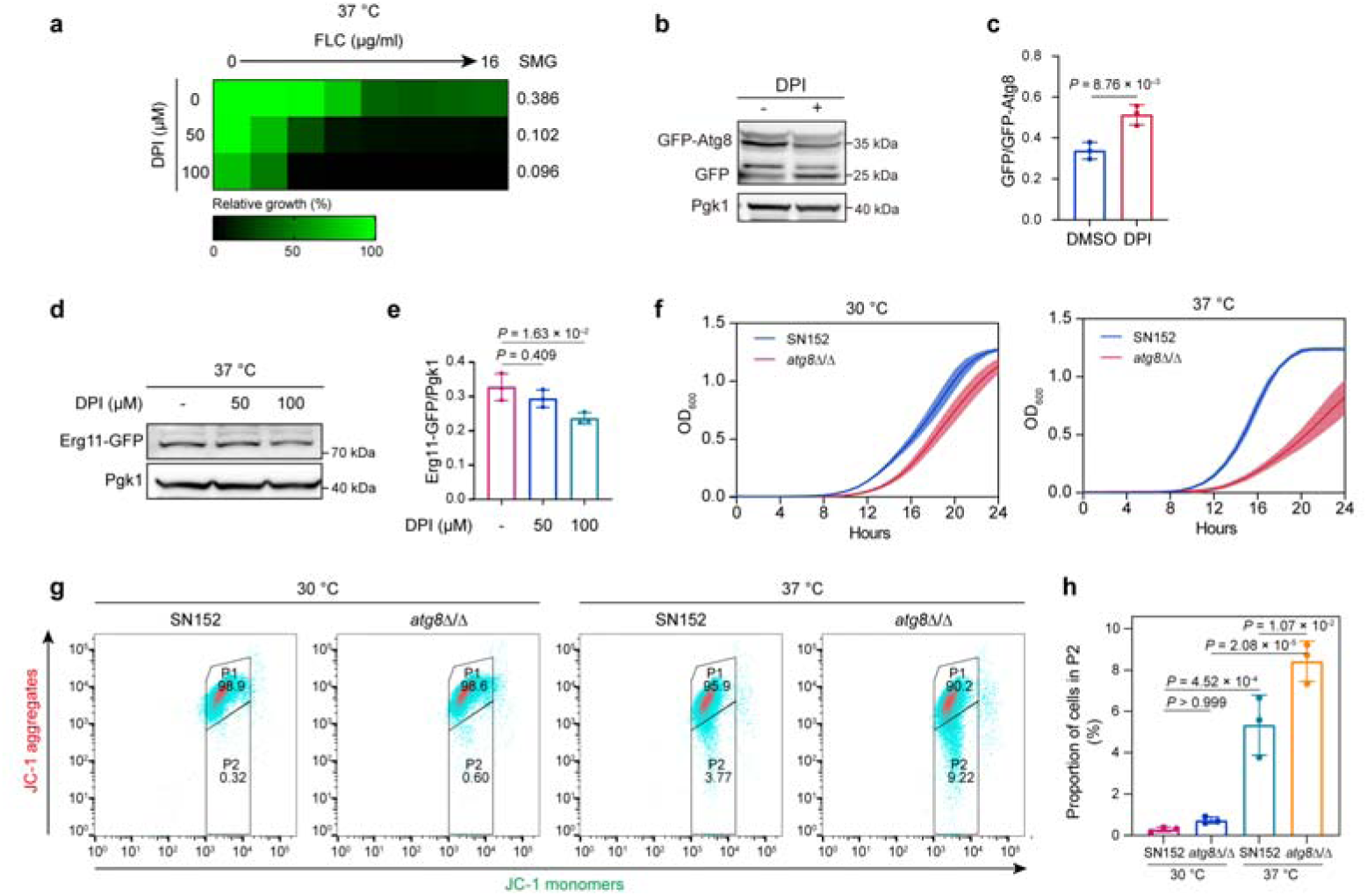
Atg8 relocalizes to mitochondria to safeguard them against oxidative damage induced by Body temperature. **a**, Broth microdilution assays showing SMG levels at 24 h for *C. albicans* SN152 cultured at 37□°C in the presence of FLC combined with the antioxidant, Diphenyleneiodonium chloride (DPI; 0 μM, 50 μM and 100 μM). Technical duplicates were averaged. **b**, GFP-ATG8 cleavage assay showing the autophagic activity after pre-cultivation with or without DPI (100 μM) at 37 °C for 4 h. **c**, Quantification of the autophagic activity displaying as the ratio of free GFP to GFP-ATG8 by Image J. Data represent means□±□s.d. of three replicates, two-tailed, unpaired t test. **d**, Degradation of Erg11-GFP in *C. albicans* SN152 in the presence of DPI (0 μM, 50 μM and 100 μM) after inhibition of protein synthesis by cycloheximide upon a 2-hour shift to 37□°C. **e**, Quantification of the degradation of Erg11-GFP calculated as the ratio of Erg11-GFP to Pgk1 by Image J. Data represent means□±□s.d. of three replicates, ordinary one-way ANOVA, Bonferroni’s multiple comparisons test. **f**, Growth curve of *C. albicans* SN152 (blue) and *atg8*Δ/Δ null mutant (red) at 30□°C (left) and 37□°C (right) for 24 h. The optical density at 600 nm (OD_600_) was measured every hour. The colored bands show the means□±□s.d. of six replicates. **g**, Determination of mitochondrial membrane potential (MMP) by JC-1 for *C. albicans* SN152 and the *atg8*Δ/Δ null mutant at 30 °C and 37 °C for 4 h through flow cytometry. JC-1 aggregates show red fluorescence (FL-1 channel, y-axis) and JC-1monomer shows green fluorescence (FL-2 channel, x-axis). Healthy cells were classified in P1, while cells with damaged mitochondria were classified in P2. The representative images shown are from three replicates. **h**, Quantification of the proportion of cells in P2 with damaged mitochondria by FlowJo. Data represent means□±□s.d. of three replicates, ordinary one-way ANOVA, Bonferroni’s multiple comparisons test.

## Discussion

In this study, we observed that short-term exposure of *C. albicans* to the human body temperature of 37□°C, as opposed to 30□°C, results in increased azole tolerance, but not resistance. This phenomenon is primarily attributed to the reduced degradation rate of Erg11 at 37□°C compared to 30□°C. The slower degradation of Erg11 at 37□°C is due to its degradation via autophagy, which is inhibited under this condition in *C. albicans*. The inhibition of autophagy at 37□°C may be mediated by elevated mitochondrial production of ROS, leading to mitochondrial dysfunction. In response to heat stress-induced mitochondrial dysfunction, autophagy-related proteins such as Atg8 accumulate around the mitochondria, while their presence in the ER diminishes, thereby suppressing autophagy in *C. albicans* (Figure 8).

**Fig. 8.**
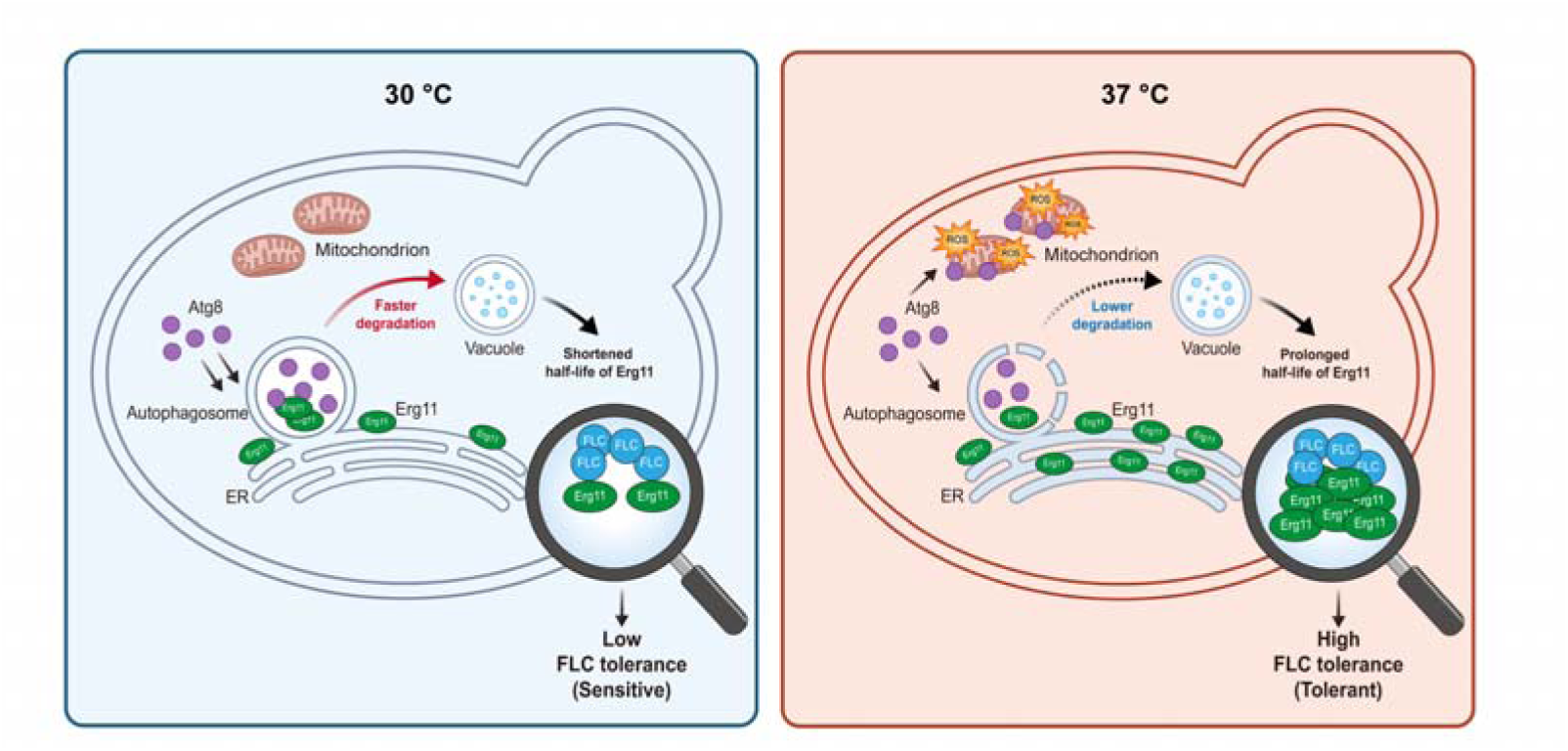
Schematic diagram showing a model of body temperature-induced azoles tolerance. Compared with 30□°C, body temperature (37□°C) results in an elevated mitochondrial production of ROS and subsequently leads to mitochondrial dysfunction which causes the relocation of Atg8 to mitochondria and the inhibition of autophagy. As the result, the slower degradation of Erg11 results in azole tolerance in *C. albicans* at 37□°C.

Our investigation revealed that short-term exposure of *C. albicans* to human body temperature markedly increases the fungus’s tolerance to azoles. This phenomenon may significantly contribute to the ineffectiveness of azole treatments in infections caused by non-resistant *C. albicans*. While the development of azole resistance in *C. albicans* is a well-established cause of therapeutic failure, azole therapies may also prove ineffective against infections involving non-resistant strains(Berman and Krysan, 2020; Feng et al., 2023). This ineffectiveness is largely due to the intrinsic tolerance of *C. albicans* to azoles(Berman and Krysan, 2020; Feng et al., 2023; Yang et al., 2023). Previous studies have confirmed that both in vitro and in vivo antifungal efficacy of FLC is substantially reduced against *C. albicans* strains exhibiting high azole tolerance(Li et al., 2024). Research has further shown that this tolerance is a pivotal factor in the persistence of candidemia(Rosenberg et al., 2018; Levinson et al., 2021). Importantly, among patients who received their initial dose of FLC within 24 h of candidemia onset, the probability of treatment failure was significantly higher when the infecting strain demonstrated elevated tolerance to FLC(Levinson et al., 2021). This study illustrates that even brief exposure to body temperature conditions can markedly increase the azole tolerance of non-resistant *C. albicans*, potentially resulting in the ineffectiveness of azole therapy for candidemia. These findings suggest that in the development of azole adjuvants to augment antifungal efficacy against *C. albicans*, research should prioritize evaluating whether these compounds facilitate the fungicidal action of azoles against *C. albicans*(Berman and Krysan, 2020; Lu et al., 2021; Xiong et al., 2025).

Erg11 is integral to the biosynthesis of ergosterol and serves as a primary target for azoles(Jordá and Puig, 2020; Van Rhijn et al., 2021). While the implications of Erg11 mutations and overexpression have been extensively explored, its stability is equally crucial for maintaining fungal susceptibility to azoles(Villasmil et al., 2020; Handelman et al., 2021; Hu et al., 2023). Current evidence suggests that Erg11 is processed via the Asi-mediated Endoplasmic Reticulum-Associated Degradation (ERAD) pathway, which involves ubiquitin-conjugating enzymes Ubc4 and Ubc7, the Asi1/Asi2/Asi3 ubiquitin ligase complexes, and Cdc48-facilitated extraction, which culminates in Erg11 ubiquitination and subsequent degradation in *S. cerevisiae*(Foresti et al., 2014; Khmelinskii et al., 2014; Natarajan et al., 2020). However, it remains uncertain whether ubiquitinated Erg11 is degraded by the 26S proteasome—a pathway commonly associated with many ERAD substrates(Krshnan et al., 2022)—or through autophagy, which is typical for membrane proteins(Li et al., 2022). In this study, we demonstrated that the degradation of Erg11 is inhibited by the autophagy inhibitor wortmannin or by the deletion of autophagy key gene *ATG8*, whereas the 26S proteasome inhibitor MG132 did not significantly affect Erg11 degradation. Additionally, we observed that Erg11 can translocate to the vacuole, suggesting that Erg11 in *C. albicans* is degraded via autophagy. Our investigation further revealed that brief exposure to a body temperature environment decelerates the degradation rate of Erg11 in *C. albicans* by inhibiting the autophagy process, thereby affecting azole drug tolerance. This finding expands the cognitive framework of azole tolerance mechanisms in *C. albicans*, highlighting that host temperature can dynamically regulate the abundance of the target enzyme Erg11 through the autophagy-dependent proteostasis regulation.

In conclusion, this study demonstrates that under physiological temperature conditions, the autophagy process in *C. albicans* is inhibited, leading to a reduced degradation rate of Erg11. This inhibition consequently enhances the tolerance of *C. albicans* to azoles. These findings suggest that promoting the degradation of Erg11 in *C. albicans* may serve as a viable strategy to augment the fungicidal efficacy of azoles in the human host.

## Methods

### Plasmids, strains, primers, agents, and cultural conditions

All plasmids, strains, and primers utilized in this study are detailed in Supplementary Tables 1, 2, and 3, respectively. The *C. albicans* strains were cultured in YPD medium, comprising 1% (w/v) yeast extract (Oxoid Ltd, Basingstoke, UK), 2% (w/v) peptone (Oxoid Ltd, Basingstoke, UK), and 2% (w/v) glucose (Sangon Biotech, Shanghai, China), at temperatures of either 30□°C or 37□°C. For the spotting assay, YPD medium plates were supplemented with 2% (w/v) agar (Sangon Biotech, Shanghai, China) and various concentrations of compounds. To construct mutant *C. albicans* strains, SD selective medium was employed, consisting of 0.67% (w/v) yeast nitrogen base without amino acids (Sangon Biotech, Shanghai, China), 2% (w/v) glucose, 2% (w/v) agar, and an appropriate amino acid mix (Sangon Biotech, Shanghai, China). FLC, itraconazole, miconazole, ketoconazole, voriconazole, fluvastatin, terbinafine, caspofungin, and amphotericin B were procured from Aladdin, Shanghai, China. Myriocin, staurosporine, brefeldin A, and amorolfine were obtained from MedChemExpress, Shanghai, China. Tunicamycin was sourced from TargetMol, Boston, USA. All these compounds were dissolved in dimethyl sulfoxide (DMSO) supplied by Sangon Biotech, Shanghai, China, and stored at -20□°C.

### Spotting assays

The spotting assay was performed as described previously (Fang et al., 2023; Li et al., 2024). FLC was dissolved in DMSO, added to YPD medium, and used to prepare solid plates. Cultures of *C. albicans* grown overnight were standardized to a concentration of 1□×□10^7^ cells/ml in YPD. These cultures were subsequently subjected to a series of 1:10 serial dilutions in YPD to five concentration gradients. The plates were incubated at either 30□°C or 37□°C for a duration of 48 h. Post-incubation, images of the plates were captured using the Tanon 5200 Imaging System (Tanon, Shanghai, China) in conjunction with AllDoc_X software.

### In vitro antifungal susceptibility testing

To examine the in vitro susceptibility to five azoles (FLC, itraconazole, miconazole, ketoconazole, and voriconazole), other ergosterol biosynthesis inhibitors (terbinafine and fluvastatin) and non-ergosterol biosynthesis antifungals (amphotericin B, caspofungin, tunicamycin, myriocin, and staurosporine), the broth microdilution assay was conducted according to previously reported protocol with slight modifications (Lu et al., 2023c). Briefly, 100 μl of drugs at 2-fold the final concentration was serially diluted in flat-bottom 96-well plates and combined with 100□μl of overnight *C. albicans* cultures adjusted to 1□×□10^3^ cells/ml. The plates were incubated separately at 30□°C and 37□°C without shaking, and the optical density at 600 nm (OD_600_) was measured with a spectrophotometer, Multiskan SkyHigh (Thermo Scientific, Singapore) after 24 h. The MIC value was determined as the drug concentration at which 50% of the growth was inhibited in terms of OD_600_ values compared to the no-drug wells.

To compare the values of SMG of FLC against different *Candida* isolates (*C. albicans*, *C. parapsilosis*, *C. glabrata*, *C. krusei*, and *C. guilliermondii*), the *C. albicans* wild-type strain SN152 and different mutants (*cdr1*Δ/Δ, *efg1*Δ/Δ, *ndt80*Δ/Δ, *upc2*Δ/Δ, *erg24*Δ/Δ, and *erg251*Δ/Δ) between 30□°C and 37□°C, based on the broth microdilution assay, the SMG value at 30□°C was calculated as the average growth per well above the MIC divided by the level of growth without the drug, and the SMG at 37□°C was calculated using the same MIC at 30 °C. The specific formula for calculating the value of the SMG is as follows:

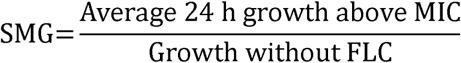

To examine whether one drug (cyclosporin A, terbinafine, fluvastatin, amorolfine, wortmannin, MG132, and NAC) combined with FLC can clear trailing in *C. albicans* at 30 °C and 37 °C, the checkerboard assays were used as described previously (Epp et al., 2010). 50 μl of 4-fold the final drug concentration of drug A was dispensed in 2-fold serial dilution steps across plate columns, and then, 50□μl of 4-fold the final drug concentration of drug B was dispensed in 2-fold serial dilution steps down rows of the plate. 100 μl of overnight *C. albicans* cultures adjusted to 1□×□10^3^ cells/ml was dispensed in all drug-containing wells plus one control well containing no drugs. Cell concentrations were recorded as the absorbance at OD_600_ after incubation at 30□°C or 37□°C for 24□h.

### The homozygous null mutants constructions

To construct homozygous null mutants in this study, *C. albicans* wild-type strain SN152 was used to permit disruption of both alleles of a given target gene with two auxotrophic markers; In this experiment, the first round of gene disruption was carried out with one of the three selectable markers, usually *Candida dubliniensis* (*C. dubliniensis*) *HIS1*, and the second round was carried out with one of the two remaining markers, usually *C. dubliniensis ARG4*. After selection of transformants on the SD medium without histidine (SD/-His) or without histidine and arginine (SD/-His/-Arg), gene disruption candidates were screened by PCR for expected 5’ and 3’ junctions as well as the size of the disrupted gene (Noble and Johnson, 2005).

### The doxycycline-inducible Tet-off promoter of genes mutants constructions

To construct the doxycycline-inducible Tet-off promoter of *ERG11* or *ENO1* mutants, first the *ERG11* and *ENO1* gene on one allele were deleted separately like what we described above to generate *ERG11*/*erg11*Δ and *ENO1*/*eno1*Δ. Then the Tet-off vectors, pCPC44 contained an *ADH1* promoter, an *ACT1* terminator and a tetracycline repressor (TetR) element between them. TetR element was codon-optimized by chemical synthesis and fused to GAL4AD to generate the fusion gene, *cat*TA. A tetracycline-responsive elements (P_Tet_) which was introduced into the downstream of selection markers (*C. dubliniensis ARG4*). Through two rounds of PCR amplification of pCPC44, the DNA cassettes with 78 bp homology regions to the target gene (P_tet-off_-*ERG11* and P_tet-off_-*ENO1*) could be transformed into *ERG11*/*erg11*Δ and *ENO1*/*eno1*Δ separately to generate the strain with the Tet-off promoter-controlled target gene. After selection of transformants on the SD medium without arginine (SD/-Arg), verification primers were used to verify the successful construction of the strain with the Tet-off promoter-controlled *ERG11* or *ENO1* (P_tet-off_-*ERG11*/*erg11*Δ and P_tet-off_-*ENO1*/*eno1*Δ)(Chang et al., 2018).

### The ectopic overexpression gene mutants constructions

To construct the ectopic overexpression gene mutants, the ectopic overexpression vectors, plasmids pCPC18-*ERG2*, pCPC18-*ERG4*, pCPC18-*ERG6*, pCPC18-*ERG9*, pCPC18-*ERG11*, pCPC18-*ERG24*, pCPC18-*ERG26* and pCPC18-*ERG27* were generated. In the plasmid pCPC18, the ADH1 promoter and the ACT1 terminator were sequentially inserted into the coding region of the *ADE2* gene and loxP-flanked selection marker, *C. maltosa LEU2*, was placed downstream of the ACT1 terminator. *ERG2*, *ERG4*, *ERG6*, *ERG9*, *ERG11*, *ERG24*, *ERG26* and *ERG27* open reading frame (ORF) fragments were PCR amplified from *C. albicans* SN152 genomic DNA and inserted into the plasmid pCPC18 backbone in one-step assembly reaction via Exo III-mediated ligation-independent cloning (LIC)(Chang et al., 2018).

The plasmids, pCPC18-*ERG2*, pCPC18-*ERG4*, pCPC18-*ERG6*, pCPC18-*ERG9*, pCPC18-*ERG11*, pCPC18-*ERG24*, pCPC18-*ERG26*, and pCPC18-*ERG27*, which we described above were used to construct the ectopic overexpression mutants of *ERG2*, *ERG4*, *ERG6*, *ERG9*, *ERG11*, *ERG24*, *ERG26* and *ERG27* in *upc2*Δ/Δ null mutant. Through two rounds of PCR amplification, the ectopic expression cassettes were transformed and integrated into the *ADE2* locus and were controlled by the constitutive *ADH1* promoter. After selection of transformants on the SD medium without leucine (SD/-Leu), verification primers were used to verify the successful construction of the strain with the ectopic overexpression gene(Chang et al., 2018).

### Galleria mellonella survival analysis

*Galleria mellonella* larvae were randomly assigned to six groups (n = 10 per group). Larvae were subjected to three treatment conditions: PBS□only controls (uninfected, no FLC treatment), infection with *C. albicans* SN152 alone, or infection with *C. albicans* SN152 plus 2□mg/kg FLC. Each condition was incubated at either 30□°C or 37□°C. Survival was assessed daily. In the infected groups, the larvae were infected with 5 μL of *C. albicans* SN152 at a concentration of 1×10^8^ cells/mL using a Hamilton syringe. The treatment groups was administered with 2 mg/kg FLC after 2 hour-infection with a single dose. Larval survival was monitored daily over a 7□day experimental period (Li et al., 2025).

### RNA isolation and qRT-PCR

Total RNA extractions from *C. albicans* and subsequent RNA-sequencing analyses were conducted in accordance with established protocols (Wang et al., 2022). RNA was isolated using the YeaStar RNA Kit (Zymo reserch, USA). The quality of RNA preparations was assessed by NanoDrop One (Thermo scientific, shanghai, China). One microgram of RNA was used for complementary DNA (cDNA) synthesis using 5 × PrimeScript RT Master Mix (Perfect Real Time) (Takara, Beijing, China). Quantitative reverse transcriptasePCR (qRT–PCR) reactions were prepared using TB Green® Premix Ex Taq™ II (Tli RNaseH Plus) (Takara, Beijing, China). cDNA mixtures were diluted 1:20 and 2 μl were used in a reaction volume of 20 μl with the primer pairs in Supplementary Table 3 with the following cycle conditions: preincubation step was 95□°C for 30s; the amplification step was 95□°C for 5s and 50□°C for 20s, repeat for 40 cycles in Bio-Rad CFX Connect Real-Time System RT-PCR 96-well (Bio-Rad, Singapore). The levels of gene expression with three replicates were determined in four separate amplifications and analyzed using the 2^-ΔΔCt method.

### RNA-seq and data analysis

*C. albicans* SN152 were overnight cultured at 30□°C and then 1:100 diluted to fresh YPD medium cultured at 30□°C and 37□°C and collected at 4-hour time point. Samples in biological triplicates were submitted to the BGISEQ platform (BGI Genomics, Wuhan, China) for RNA sequencing analyzes. Transcriptome libraries were generated using the Standard Sensitivity RNA Analysis Kit (15 nt) (DNF-471) following the manufacturer’s protocols performed by Fragment Analyzer. The raw RNA-seq reads were quality controlled with SOAPnuke (v1.5.6)(Li et al., 2008). The trimmed reads were mapped to the final assembly genomes *C. albicans* SC5314 (reference candidagenome: Assembly22) using HISAT2 (v2.1.0)(Kim et al., 2015). Bowtie2 (v2.3.4.3)(Langmead and Salzberg, 2012) was used to compare clean reads to reference gene sequences, and then RSEM (v1.3.1)(Li and Dewey, 2011) was used to calculate gene expression levels for individual samples. DESeq2 (v1.4.5)(Love et al., 2014) was used to perform differential gene testing under the condition of Q value ≤ 0.05 or FDR ≤ 0.001. Kyoto Encyclopedia of Genes and Genomes (KEGG) and GO enrichment (GO) enrichment analysis further explored the gene functions associated with phenotypic changes. Based on hypergeometric testing, we used Phyper to perform GO on differentially expressed genes enrichment analysis. The gene with Qvalue ≤ 0.05 was defined as significant enrichment in candidate genes that meet this condition. The value of log2 Fold change (37□°C vs 30□°C) was not directly calculated by taking the mean expression level (FPKM) and then calculating the log. Instead, it was standardized using the local standard deviation of the FPKM (counts) based on a linear regression of all gene expression levels, taking a small interval of the corresponding gene, and then calculating the corresponding standard deviation. When correcting for each mRNA expression level, it was related to the distribution of all mRNA expression levels in each sample. Gene set enrichment analysis was performed with the GSEA software(Subramanian et al., 2005). Heatmaps were generated using the pheatmap R package, with clustering distance and method set to Euclidean and ward.D2, respectively (Tuling Biotechnology, Shanghai, China).

### Protein extractions and western blotting

The protein extractions and western blotting assay were conducted following a previously established protocol with minor modifications (Fang et al., 2025). *C. albicans* cells were harvested at 4 °C, resuspended in 500 μl of protein lysis buffer which consists of phosphate-buffered saline (PBS; Sangon biotech, Shanghai, China), 5 mM EDTA (pH 8.0; Sangon biotech, Shanghai, China), 1mM PMSF (Sangon biotech, Shanghai, China), and 1.0% Protease Inhibitor cocktail (TargetMol, Shanghai, China), transferred into screw-cap tubes with the addition of 500 μl of glass beads, and homogenized by a bead-beater for 4 cycles of 1min, with 1 min cooling on ice in each cycle. Then the tubes were centrifuged at 10,000 × g for 10 min at 4□°C and supernatant was transferred to a new tube. The protein concentration was measured with Bradford Protein Assay Kit (Epizyme, Shanghai, China). Equal amounts of protein were separated by 8-20% SDS-PAGE (Byotime, Shanghai, China) and then transferred onto PVDF membranes (Merck Millipore Ltd., Carrigtwohill, Ireland).

After blocking with 1 ×Protein Free Rapid Blocking Buffer (Epizyme, Shanghai, China) for 10 min at room temperature, the membranes were incubated with primary antibodies against GFP (Anti-GFP [B-2]) mouse monoclonal antibody, 1:1000 dilution; Santa Cruz Biotechnology, USA) or α Tubulin (Anti-α Tubulin [YL1/2] rat monoclonal antibody, 1:1000 dilution; Santa Cruz Biotechnology, USA) or Pgk1 (Anti-Pgk1 [14] mouse monoclonal antibody, 1:1000 dilution; Santa Cruz Biotechnology, USA) overnight at 4□°C. Then, the membranes were incubated with secondary antibodies (Peroxidase-conjugated Anti-Mouse IgG_k_, 1:10000 dilution; Santa Cruz Biotechnology, USA) for 1 h at room temperature. The probed protein was visualized using an ECL western blotting kit (Epizyme, Shanghai, China) and photographed by the Tanon 5200 Chemiluminescence Imaging System (Tanon, Shanghai, China) with AllDoc_X software. The α Tubulin and Pgk1 were used as the loading control.

### Laser scanning confocal microscopy

The laser scanning confocal microscopy assay was conducted as previously described (Zhen et al., 2024a). For the colocalization of Upc2 and nucleus, the Upc2-GFP mutants cultured overnight at 30□°C were 1:10 diluted to YPD medium containing 16 μg/ml FLC and cultured at 30□°C or 37□°C for 4 h. After washing twice with PBS, cells were collected and incubated with 10 μg/ml the nuclear dye, Hoechst 33342 (MedChemExpress, Shanghai, China) for 10 min at room temperature in the dark. Finally, cells were washed with PBS twice and photographed. The fluorescence distribution curve of the GFP green fluorescence and nuclear dye (Hoechst) with a red fluorescent pseudocolor in the region of interest in single cell was used to show the colocalization of Upc2 and nucleus at 30□°C and 37□°C by Image J (1.54g).

For the colocalization of Erg11 and vacuole, the Erg11-GFP mutant grown overnight were 1:10 diluted to YPD medium and cultured at 30□°C or 37□°C for 1 h. After washing twice with PBS, cells were collected and incubated with 100 μM vacuole dye, CellTracker Blue CMAC (MedChemExpress, Shanghai, China) for 30 min at 30□°C or 37□°C in the dark. After washing with PBS twice, the cells were photographed. The colocalization rate was calculated by dividing the number of cells colocalized showing yellow fluorescence signal by the total number of cells.

For comparing the autophagy activity between 30□°C and 37□°C, the GFP-Atg8 mutants grown overnight were 1:10 diluted to YPD medium and cultured at 30□°C or 37□°C for 1 h. After washing twice with PBS the cells were photographed. The autophagy activity is reflected as the ratio of the number of cells with GFP-ATG8 dots to the total number of cells at 30□°C and 37□°C.

For the colocalization of Atg8 and mitochondria or ER, the GFP-Atg8 mutants grown overnight were 1:10 diluted to YPD medium and cultured at 30□°C or 37□°C for 30 min. After washing twice with PBS, cells were collected and incubated with 0.3 μM mitochondria dye, MitoTracker Deep Red FM (MedChemExpress, Shanghai, China) or 1 μM ER dye, ER-tracker Red (MedChemExpress, Shanghai, China) for 30 min at 30□°C or 37□°C in the dark. After washing with PBS twice, the cells were photographed. The colocalization ratio was calculated by dividing the number of GFP-Atg8 dots colocalized with mitochondria or ER by the total number of cells containing GFP-Atg8 dots.

All the images were acquired on an Andor Dragonfly Confocal Imaging System (Andor, Oxford instrument, UK), ImarisViewer (10.1.0) equipped with a white light laser. A 100 ×□oil-immersion objective was employed. The excitation (Ex)/emission (Em) wavelength for GFP was 488□nm/521 nm. The Ex/Em wavelength of Hoechst 33342 for nucleus was 405 nm/445 nm. The Ex/Em wavelength of CellTracker Blue CMAC for vacuole was 405 nm/445 nm. The Ex/Em of MitoTracker Deep Red FM for mitochondria was 637 nm/660 nm. The Ex/Em of ER-tracker Red for ER was 572 nm/582 nm□-□750 nm.

### Quantitation of the content of sterols

The content of sterols in *C. albicans* was determined by a method described previously with slight modifications (Lu et al., 2023b; Hang et al., 2025). *C. albicans* SN152 were overnight cultured at 30□°C and then 1:100 diluted to fresh YPD medium cultured at 30□°C or 37□°C and collected at 4-hour, 8-hour and 16-hour time points. For the group with FLC, *C. albicans* SN152 were overnight cultured at 30□°C and then 1:100 diluted to fresh YPD medium with 16 μg/ml FLC, cultured at 30□°C and 37□°C for 16 h. Cells were collected and washed three times with double-distilled water (ddH_2_O), and the weight of the wet cell pellet was adjusted to approximately 0.5 g. Furthermore, the 15% (m/v) NaOH (Sangon biotech, Shanghai, China) dissolved in 90% (v/v) ethanol (Sangon biotech, Shanghai, China) were added to each pellet and mixed thoroughly. The suspension was incubated at 80 °C for 1 h in a water bath. Sterols were extracted by adding 18□ml of petroleum ether (boiling range: 30□°C-60□°C; SCR, Shanghai, China) in total three times, and each time the mixture was vortexed vigorously for 2□min and allowed to stand for 3□min. The petroleum ether layer was transferred to the same clear glass tubes and washed twice with ddH_2_O. The collected petroleum ether was volatilized at 65□°C for 20□min in the water bath. Finally, the sterols were extracted by the addition of 600□μl of n-hexane (Sangon biotech, Shanghai, China), and a 200-μl aliquot was injected for gas chromatography-mass spectrometry (Finnigan Voyager, USA) with HP-50 columns (50% phenyl–50% methylpolysiloxane, 30 m by 0.25□mm by 0.25□μm). Before injection, 20□mg of cholesterol (Sangon biotech, Shanghai, China) was added to each sample as the internal reference for quantifying other sterols. Sterols of interest were identified by their relative retention times, and mass spectra were compared with the sterol profiles of NIST. The contents of sterol were calculated as the sterol mass corresponding to the unit mass of cells (μg/mg). Assays were performed in duplicate three times.

### Cycloheximide chase analysis

Cycloheximide (CHX) chase analysis was essentially performed by a method with some modifications (Buchanan et al., 2016; Hang et al., 2025). Briefly, The Erg11-GFP mutants were inoculated overnight in YPD at 30□°C with shaking. The overnighted cultures were diluted to an OD_600_ value of 0.2 in fresh YPD medium and incubated at 30□°C with shaking for about 4 h, until the cells reach mid-logarithmic growth phase (OD_600_=1.0). Then 25 OD_600_ units of each culture per time point were collected. Then each cell pellet was resuspended in 10 ml of 30□°C or 37□°C pre-warmed fresh YPD medium per 25 OD_600_ units of cells. After the equilibrium of cell suspensions by incubation for 5 min in the 30□°C or 37□°C heat block, cycloheximide (MedChemExpress, Shanghai, China) was added to a final concentration of 1 mg/ml to the cell suspension and the timer started counting from 0:00. Immediately after adding cycloheximide and vertexing, 10 ml (∼25 OD_600_ units) of the cell suspension with added cycloheximide was transferred to a pre-cooling microcentrifuge tube to centrifuge at 4,000 × g at 4□°C for 5 min. After that, the cell suspension was removed the supernatant and washed with sterile water twice at 4,000× g at 4□°C for 5 min and stored at -20□°C. When all samples at different time points have been collected, the cells are ready to lysis for western blotting like we described above. To investigate the effects of Wortmannin, MG132 and Diphenyleneiodonium chloride (DPI) on the degradation of Erg11, Wortmannin (1 μM), MG132 (100 μM) and DPI (100 μ M) were added after adding cycloheximide to inhibit the protein synthesis. Pgk1 was used as the loading control. Assays were performed in duplicate three times.

### GFP–ATG8 cleavage assay

The GFP–ATG8 cleavage assay was conducted as previously described(Zhen et al., 2024a). To investigate the effect of different temperatures on the activity of autophagy, the GFP-Atg8 mutants were overnight cultured and then 1:10 diluted in YPD medium at 30□°C or 37□°C for 4 h. And investigate the effect of DPI on the activity of autophagy, the GFP-Atg8 mutants were overnight cultured and then 1:10 diluted in YPD medium at 37□°C in the presence of 100 μM DPI for 4 h. Total proteins were extracted, measured with Bradford Protein Assay Kit and immunoblotted with primary antibodies against GFP like what we describe in the western blotting assay. Pgk1 was used as the loading control. The GFP cleavage activity was quantified by determining the ratio of free GFP to GFP–ATG8. Bafilomycin A1 inhibition assays To monitor autophagic flux in *C. albicans*, the GFP□Atg8 reporter mutants combined with pharmacological inhibition of vacuolar degradation using bafilomycin A1 (Baf A1, MedChemExpress, Shanghai, China) were employed. In this assay, the GFP-Atg8 mutants were overnight cultured and then 1:10 diluted in YPD medium at 30□°C or 37□°C. 2 μM Baf A1 was added after an initial 2-h culture period under the indicated temperatures, with a treatment duration of 2 h. Total proteins were extracted, measured with Bradford Protein Assay Kit and immunoblotted with primary antibodies against GFP like what we describe in the western blotting assay. Pgk1 was used as the loading control. The accumulation of GFP-Atg8 in Baf A1□treated samples indicates the rate of autophagosome synthesis, because vacuolar degradation is blocked. For each sample, the Baf A1□induced accumulation of full-length GFP-Atg8 at 30 °C or 37 °C was determined by subtracting the normalized signal intensity of the vehicle□treated (without BafA1) sample from that of the Baf A1□treated sample (Δ = Baf A1-vehicle) (Paumier et al., 2025). This value represents the rate of autophagosome synthesis during the 2□h Baf A1 treatment period.

### Measurement of intracellular reactive oxygen species

Intracellular ROS was measured using 2’,7’-dichlorodihydrofluoroscein diacetate (DCFH-DA) with some modifications (Moraitis and Curran, 2004; Li et al., 2024). *C. albicans* SN152 were overnight cultured at 30□°C and then 1:5 diluted and treated with 10 μM DCFH-DA (MedChemExpress, Shanghai, China) and incubated at 30□°C or 37□°C with shaking for 30 min. The cells were harvested at 4□°C and washed twice with PBS. Then cells were suspended in 500 μl PBS and transferred into screw-cap tubes with the addition of glass beads equal in volume to the cell pellet, and homogenized by a bead-beater for 4 cycles of 1 min, with 1 min cooling on ice in each cycle. Then the tubes were centrifuged at 10,000 × g for 10 min at 4 °C and supernatant was transferred to a new tube. The fluorescence was recorded, at 25 °C, at an excitation wavelength (Ex) of 485 nm and an emission (Em) wavelength of 535 nm, using the microplate reader, Infinite F Plex (Tecan, Groedig, Austria). Em measurements were normalized by estimation of the protein content of the supernatant using a Bradford protein assay kit. The SFI was expressed as fluorescence [(Em = 535 nm/Ex = 485 nm)/mg protein]. As a control, background fluorescence of cells not treated with DCFH-DA, was also recorded. In order to ensure that the fluorescence readings were in the linear dynamic range of DCF concentrations (1 nM – 400 nM), a calibration curve was constructed of fluorescence (Em = 535 nm, Ex = 485 nm) vs. DCF concentration.

### Growth curve assays

Growth curve assay was performed as described previously (Lu et al., 2015; Wu et al., 2024). *C. albicans* SN152 and *atg8*Δ/Δ null mutant were overnight cultured at 30□°C. Then cells were adjusted to the concentration of 10^3^ cells/ml by the hemacytometer. Then cells were cultured at 30□°C or 37□°C with shaking (200 rpm). The optical density at 600 nm (OD_600_) was measured every hour until the stationary phase of the growth curve was reached by using the microplate reader, Infinite M Nano (Tecan, Groedig, Austria).

### Mitochondrial Membrane Potential (MMP) assay

Changes in MMP were determined by the MMP probe JC-1 as previously described with some modification (Zhao et al., 2025). *C. albicans* SN152 and *atg8*Δ/Δ null mutant were overnight cultured at 30 °C and then 1:10 diluted to fresh YPD medium cultured at 30□°C or 37□°C for 4 h. Then cells were washed and resuspended in PBS with 2.5 μg/ml JC-1 at 37□°C for 20 min in the dark, then washed twice. All samples were estimated via the BD LSRFortessa Cell Analyzer (LSRFortessa; BD Bioscience, USA). Gating of single cells was achieved as follows. The green fluorescence signal of the JC-1 monomer was detected using the FITC channel equipped with a 530□nm band-pass filter (bandwidth 30□nm) while the red fluorescence signal of JC-1 aggregates was detected using the PE channel equipped with a 585□nm band-pass filter (bandwidth 15□nm). First, the cells populations were separated from cell debris and selected via SSC-A (side scatter; U = 302 V) and FSC-A (forward scatter; U = 640 V) signals. Second, duplet cells were excluded via FSC-W and FSC-H signals. A total of 50,000 events was analyzed per sample. Flow cytometry data were collected using FACS Diva Software and analysed using FlowJo (10.8.1). SN152 cells cultured at 30 °C for 4 h with healthy mitochondria were considered as negative control sample. Here, according to the negative control sample, P1 was set as a region in which > 95% of the dual fluorescent population falls within it and the JC-1 red/green fluorescence ratio > 1(Guthrie and Welch, 2006; Yamakawa et al., 2013; Elefantova et al., 2018). The loss of MMP was reflected by the proportion of cells in P2 with less aggregates (red fluorescence) and more monomer (green fluorescence) among different groups. The ordinary one-way ANOVA, Bonferroni’s multiple comparisons test was used to evaluate multiple comparisons of the proportion of cells with damaged mitochondria.

### Statistical analysis

Statistical analysis was performed using Graphpad Prism 9.0 software. The two-tailed paired t-test was used to compare the values of SMG of FLC against clinical *Candida* isolates (*C. glabrata*, *C. krusei* and *C. guilliermondii*) between 30 °C and 37 °C. The Wilcoxon matched-pairs signed rank test was used to compare the values of SMG of FLC against *C. albicans* and *C. parapsilosis* between 30 °C and 37 °C. The two-tailed unpaired t-test was used to compare the expression of proteins in WB, sterol content, the degradation of Erg11-GFP, the colocalization rate, the autophagic activity, Atg8 dots and ROS level between two groups. The ordinary one-way ANOVA, Bonferroni’s multiple comparisons test was used to evaluate multiple comparisons of the degradation of Erg11-GFP in the presence of different compounds and the proportion of cells with damaged mitochondria. For all analyses, *P*□<□0.05 was considered significant. Data are shown as means□±□standard deviation (s.d.) from three or more replicates.

## Data availability

The RNA sequencing data of *C. albicans* SN152 cultured at 30 °C and 37 °C for 4 h are publicly available at the NCBI Sequence Read Archive (SRA) and can be accessed under BioProject PRJNA994182.

## Acknowledgments

This study received financial support from the National Natural Science Foundation of China (No. 82574464) and the Innovation Program of Shanghai Municipal Education Commission (202101070007-E00094). We extend our heartfelt thanks to Jianpeng Zhang and Xiaochao Zhao for their professional support and insightful contributions throughout the data analysis process.

## Author Contributions

**YR.F.** conducted most of the experiments and performed data analysis. **YR.F.**, **H. L., M.W.,** and **YY.J.** wrote the manuscript draft and revised the manuscript. **YR.F., W.L., C.Z., XY.F,** and **XQ.S.** constructed mutant strains. **YR.F., W.L., C.Z., XY.F,** and **XQ.S.** evaluated the antifungal activities and data collection. **YR.F., H. L., M. W.,** and **YY.J.** discussed and analyzed the data. **H.L.** and **YY.J.** conceived the idea. **H.L.** and **YY.J.** directed the experiments.

